# Architectural fragility of gene regulatory networks underlies hematopoietic stem cell aging

**DOI:** 10.64898/2026.06.03.729976

**Authors:** Harlan P. Stevens, Ali Doğa Yücel, Russell A. Gould, George Cai, Vadim N. Gladyshev, Alexandru M. Plesa, George M. Church

## Abstract

Hematopoietic stem and progenitor cell (HSPC) aging contributes to immune dysfunction and age-associated disease, but its regulatory mechanisms remain unclear. Here, we present the largest single-cell multiome atlas of human circulating HSPCs to date, with >380,000 paired RNA and ATAC profiles across 77 donors. Beyond recapitulating established hallmarks of HSPC aging, we reconstructed a high-resolution gene regulatory network and identified a global rewiring in which stress-response and myeloid transcription factor (TF) programs expand, while self-renewal and lymphoid lineage-defining circuitry collapses. We associate these phenotypes with increased cis-regulatory entropy, including elevated transcriptomic noise, weakened peak-to-gene coupling, and chromatin peak broadening. This rewiring is selective, where TFs with GC-rich, promoter-proximal architectures are preserved or amplified, and complex, distal enhancer-dependent identity networks are eroded. Thus, these findings suggest that progressive entropic destabilization of gene regulatory architecture simultaneously drives stress hyperactivation, myeloid bias, and identity loss in aging HSPCs.

---

Aging is one of the most fundamental unsolved problems in biology, and it still lacks a mechanistic understanding, and is broadly defined by its effects – a progressive functional decline and elevated disease incidence and mortality. With nearly every cell in the human body residing within 50-100 μm away from a blood vessel^1^, the hematopoietic system is a central regulator of organismal health and longevity. Consequently, hematopoietic deterioration may act as a systemic amplifier of age-related pathology, manifesting as chronic inflammation, immunosenescence, anemia, increased clonal hematopoiesis of indeterminate potential (CHIP), and myeloid malignancies^2–6^. These cascading dysfunctions ultimately trace back to hematopoietic stem and progenitor cells (HSPCs), which sustain all blood and immune lineages. Thus, age-related HSPC dysfunction is both a driver of hematologic disease and a high leverage point for intervention in aging.

Tracing these organism-wide phenotypes to the cellular level, several studies have demonstrated that aged HSPCs undergo a loss of self-renewal capacity, an expansion of the phenotypic stem cell pool, and a skewed differentiation trajectory toward the myeloid lineage^7,8^. While this age-associated “myeloid bias” was traditionally viewed as a uniform lineage shift, recent high-resolution clonal tracking and single-cell analyses support a more nuanced model: a disproportionate loss of lymphoid potential accompanied by clonal compensation within the myeloid compartment^9–11^. Moreover, while cell-extrinsic factors such as chronic inflammatory remodeling of the bone marrow niche^12^ are recognized as upstream triggers, emerging evidence suggests that the resulting functional decline is ultimately locked in by profound, cell-intrinsic epigenetic and transcriptional rewiring^13–16^. Here, we use cell-intrinsic aging to refer to age-associated molecular changes observed within phenotypically matched HSPC populations, after accounting for shifts in cell-type composition and measured donor covariates.

Previous multi-omics studies have mapped these epigenetic shifts, consistently identifying the hyperactivation of stress-response pathways, particularly those driven by the AP-1 transcription factor (TF) complex^17,18^. Recently, the “SIPHON” model proposed a mechanism for this phenomenon, suggesting that AP-1 overactivation hijacks essential transcriptional co-factors, leading to the closing of chromatin at cell identity-associated loci^19^. Additionally, AP-1 activation has been shown to induce persistent epigenetic states that prime HSPCs for dysfunction and are inherited intrinsically across lineages^20,21^. While the SIPHON model alone partially explains the selective vulnerabilities of aging HSPCs, it remains unclear why AP-1 activation would preferentially degrade self-renewal and lymphoid programs while sparing, or even enhancing, myeloid trajectories, which themselves rely on lineage-specific TFs. Thus, a comprehensive, unbiased understanding of how aging reshapes gene regulatory networks (GRNs) and induces subsequent phenotypic decline is still lacking^22^.

To address this gap, we used single-cell multiome (paired scRNA-seq and scATAC-seq)^23^ profiling to map the coupled epigenetic and transcriptomic states of human circulating HSPCs across adulthood. This multimodal approach enabled reconstruction of age-associated GRNs^24^ at the level of individual regulons, defined here as transcription factors together with their inferred target genes. This allowed us to quantify network rewiring and TF activity changes with single-cell resolution. With this framework, we confirmed a robust, age-dependent increase in stress-related (AP-1) TF activity coupled with a decline in identity programs^19,25^. We also found an increase in myeloid and erythroid TF activity, coupled with a reciprocal global decline in the core programs governing lymphoid differentiation and HSC and self-renewal.

Integrating these findings, we found that the TFs preserved or amplified with age share distinct regulatory architectures from those that decline, suggesting that the structural properties of a regulon, not its lineage role alone, shape its vulnerability to aging. We propose that hematopoietic aging is driven by an entropic destabilization of regulatory architecture, in which complex, distal regulatory programs are progressively eroded while robust, proximal programs are preserved or expanded.

## Results

### Single-cell multiome atlas of human circulating HSPCs captures the hallmarks of hematopoietic aging

To map the cell-intrinsic changes in human hematopoietic aging, we generated, to our knowledge, the largest paired single-cell multiome atlas of circulating CD34+ hematopoietic stem and progenitor cells (HSPCs) to date, spanning 77 healthy donors aged 17–81 years (Fig. 1a). PBMCs were isolated from leukocyte reduction system chambers and enriched for CD34+ cells. After quality control, we retained 385,509 high-quality paired RNA and chromatin accessibility profiles.

**Figure 1:**
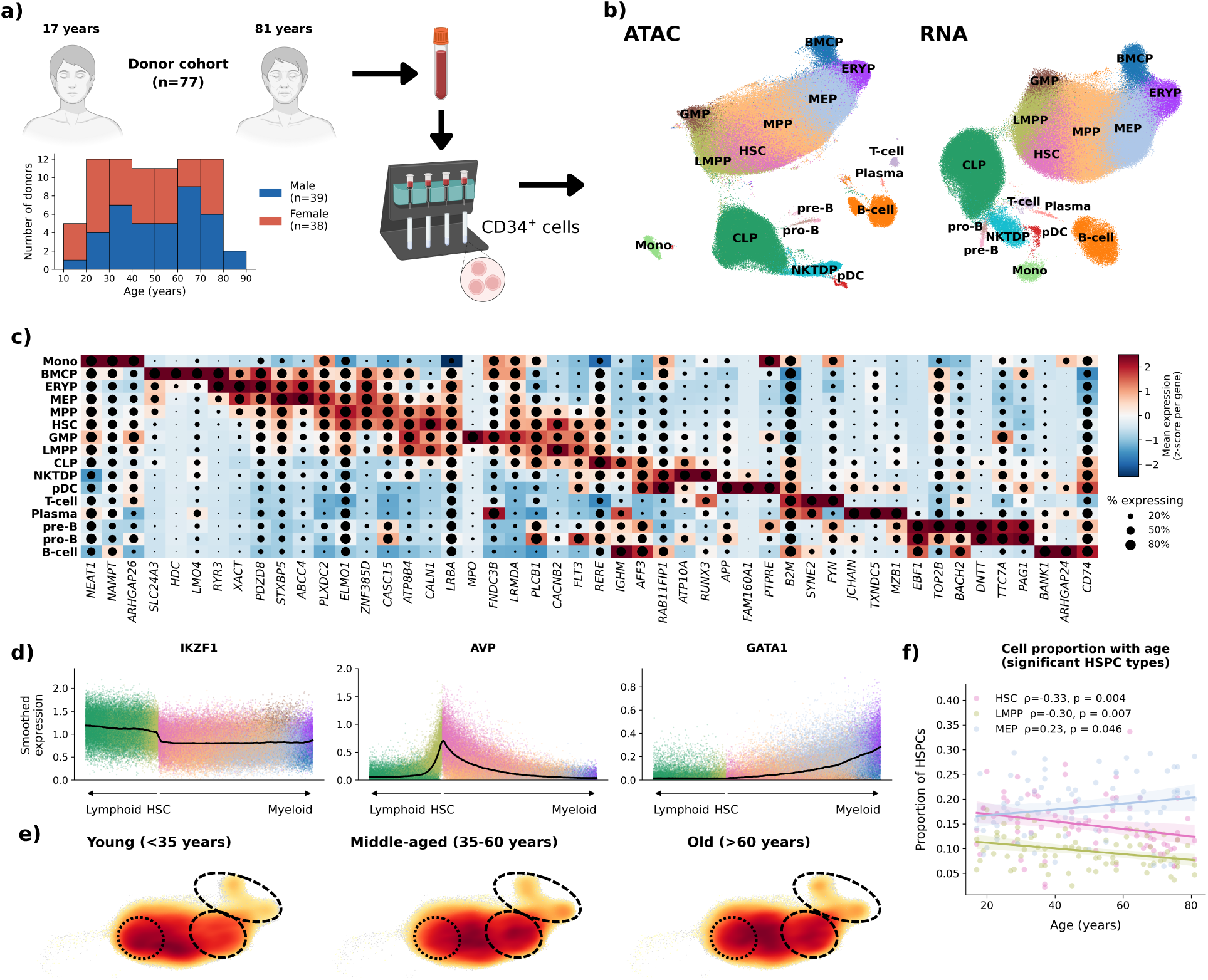
Single-cell multiome atlas of circulating human HSPCs. **a)** Schematic overview of study design with single-cell multiome profiling of circulating CD34^+^ hematopoietic stem and progenitor cells (HSPCs) across human aging. **b)** Annotated two-dimensional UMAP embedding of high-quality cells, colored by cell type. RNA UMAP shown on the right, and ATAC UMAP on the left. **c)** Top three marker genes for each annotated cell type, ranked by log fold change relative to all other cell types. **d)** Example TF expression dynamics across the lymphoid–HSC–myeloid axis. Expression was averaged over nearest neighbors and cells were ordered by HLF expression, with lymphoid-lineage cells shown on the left and HSCs and myeloid-lineage cells shown on the right. Each panel displays smoothed average log_2_ expression for one TF. **e)** Density of LMPP and myeloid-HSPC cell states across donor age groups, showing age-associated shifts in cellular composition. Red indicates the highest cell density. **f)** Cell-type proportions (relative to the myeloid-HSPC compartment) versus donor age, for HSPC populations significantly associated with age (Spearman ρ, p < 0.05). Lines show LOWESS fits.

The atlas showed high quality across both modalities, with strong cell-level RNA and ATAC metrics, high *CD34* expression across the progenitor compartment, and minimal evidence of age-associated technical confounding (Extended Fig. 1a-d). We observed a slight age-related decrease in recovered HSPCs, consistent with prior reports and likely biological in origin^26,27^. Because male donors were older than female donors in our cohort (median 57 vs. 43 years, p = 0.03; Extended Fig. 1e-f), sex was included as a covariate in all age-association analyses.

Our cell type annotation, based on clustering, marker genes, and external reference mapping, resolved the expected early hematopoietic hierarchy^10^. We recovered the HSPC cell types: hematopoietic stem cells (HSCs), multipotent progenitors (MPPs), lymphoid-primed multipotent progenitors (LMPPs), common lymphoid progenitors (CLPs), megakaryocyte-erythroid progenitors (MEPs), erythroid progenitors (ERYPs), granulocyte-monocyte progenitors (GMPs), basophil/mast cell progenitors (BMCPs), and natural killer/T-cell/dendritic cell progenitors (NKTDPs). We also captured a minor fraction of more differentiated, low-CD34^+^ populations, including plasma cells, B-cells and their precursors (pro-B and pre-B), T-cells, plasmacytoid dendritic cells (pDCs), and monocytes (Mono) (Fig. 1b-c, Extended Fig. 1b). Cell states showed strong concordance across external datasets^28–30^, confirming the accuracy of our annotations (Extended Fig. 2a-d).

The overall manifold recapitulated canonical human hematopoiesis^31^, with a central HSC node connected to downstream progenitors and a bifurcating myeloid arm (Fig. 1b). Marker genes and lineage-defining TF gradients matched expected lineage structure across the lymphoid, primitive, and myeloid compartments (Fig. 1c-d). Because circulating HSPCs can differ subtly from marrow-resident counterparts, primarily in quiescence^32^, we projected our dataset onto a matched reference containing both circulating and marrow-derived HSPCs^27^. As expected, our cells aligned most strongly with the peripheral CD34^+^ compartment and had lower cell-cycle activity than marrow-derived HSPCs (Extended Fig. 2e-i), although cycling activity was positively correlated with age (Extended Fig. 2j).

Leveraging the continuous age distribution of the cohort, we observed progressive remodeling of the HSPC compartment across adulthood, with expansion of myeloid-biased progenitors, particularly MEPs, and decline of primitive and lymphoid-associated compartments, including HSCs and LMPPs (Fig. 1e-f). We also found reduced B cells and B-cell precursors with age (Extended Fig. 3a-c), consistent with diminished lymphoid output^33^, although CD34^+^ enrichment limits the interpretation of changes in differentiated cell types. These patterns were consistent in a complementary, cluster-agnostic differential abundance analysis^34^ (Extended Fig. 3d-e).

These analyses establish our circulating HSPC multiome atlas as a representative model of human hematopoiesis that captures the major compositional hallmarks of hematopoietic aging and provides a foundation for dissecting the underlying regulatory changes.

### A shared transcriptomic aging signature across HSPCs

To define the transcriptional signature of HSPC aging, we generated donor-level pseudobulked profiles of all HSPCs and performed differential expression analysis, modeling age as a continuous variable while controlling for sex. This identified 1,004 age-associated genes (FDR < 0.05), including 594 that increased and 410 that decreased with age (Fig. 2a).

**Figure 2:**
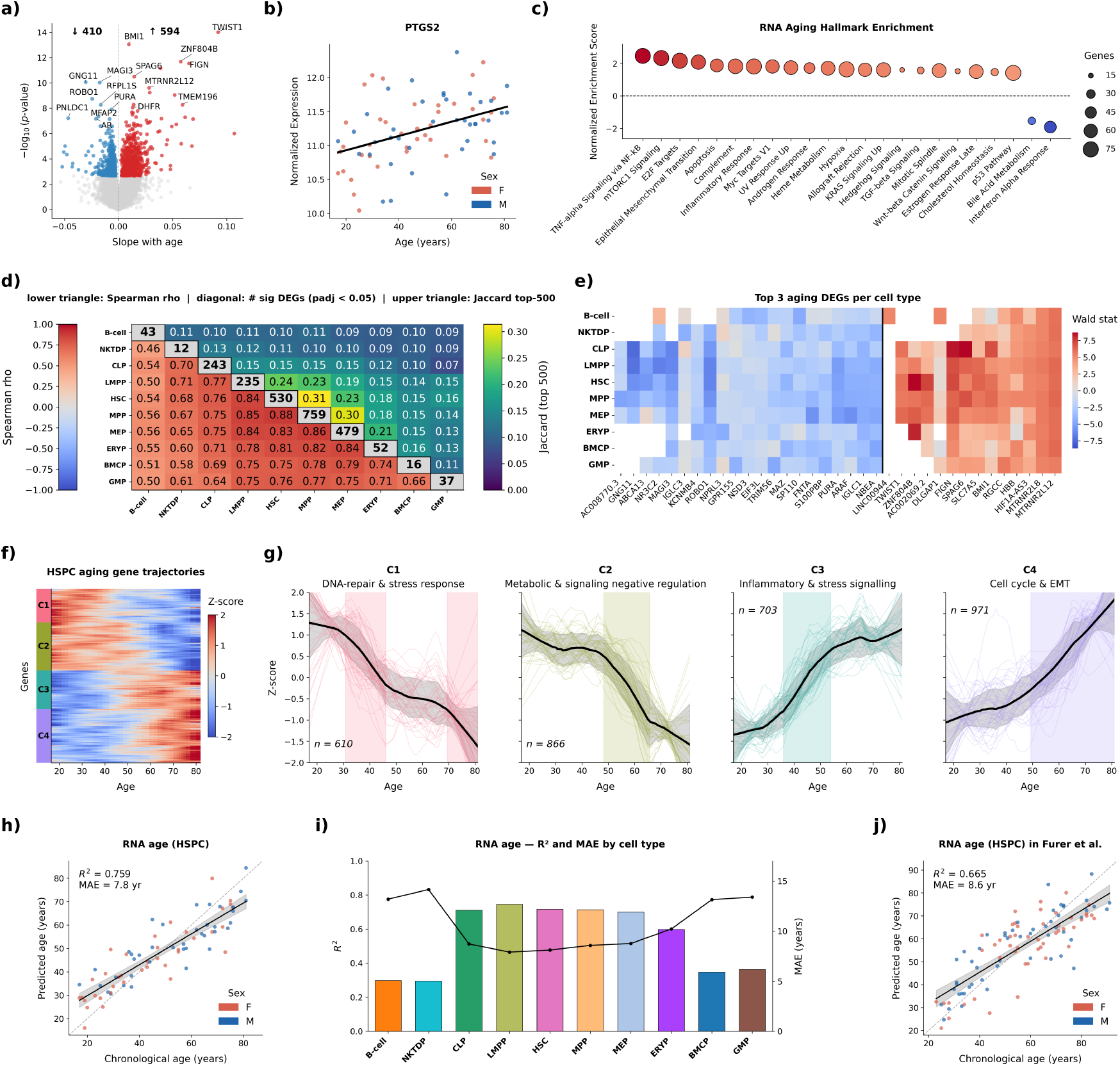
Transcriptomic signatures of HSPC aging. **a)** Volcano plot of age-associated genes in HSPCs. The x-axis shows the expression slope with age, calculated by pseudobulking all HSPCs by donor and fitting a DESeq2 model with age as a continuous variable and sex as a covariate. Using an FDR cutoff of 0.05, we identified 1,004 differentially expressed genes, with 594 increasing and 410 decreasing with age. **b)** Expression of the example age-associated gene PTGS2 across donors, plotted against donor age and colored by sex. Points show variance-stabilized pseudobulk expression in HSPCs. **c)** Gene set enrichment analysis of the HSPC aging signature. Gene sets are ordered by normalized enrichment score; dot size indicates gene set size and color indicates FDR. **d)** Similarity of transcriptomic aging signatures across HSPC cell types. Diagonal values indicate the number of differentially expressed genes (FDR < 0.05) identified within each cell type. The lower triangle shows Spearman correlations of Wald statistics across the cell type pairwise union of the top 500 most age-associated genes, and the upper triangle shows the Jaccard overlap between the corresponding top 500 gene sets. **e)** Cross-cell-type overlap of the strongest age-associated genes. Shown are the union of the top three genes increasing and top three genes decreasing with age in each cell type, colored by Wald statistic. White indicates expression too low for reliable testing. **f)** Aging trajectories of age-associated genes in HSPCs. Heatmap shows smoothed, z-score-normalized expression across donor age for genes significantly associated with age (dCor FDR < 0.05; see Methods). **g)** Unsupervised clustering of age-associated gene trajectories identified four major expression patterns with distinct temporal dynamics and associated GO term enrichments (Extended Fig. 4g-h). Black lines indicate the mean trajectory for each cluster, and colored lines show individual gene trajectories. The grey envelope around the mean denotes ±1 SD across genes in the cluster, and colored vertical shading marks age intervals over which the mean trajectory’s slope exceeds 0.05 z-score/year. **h)** Predicted versus chronological age for the transcriptomic HSPC aging predictor under leave-one-donor-out cross-validation. **i)** Transcriptomic aging predictor performance by cell type. Bar height indicates R^2^, and overlaid lines indicate MAE. **j)** Age prediction performance of the HSPC transcriptomic aging predictor applied to the Furer et al. reference HSPC atlas.

Age-upregulated genes were enriched for inflammatory, stress-response, and EMT-related programs, consistent with the well-described inflammatory remodeling of aged HSPCs^35,36^ and exemplified by the progressive increase of the canonical inflammatory/stress gene *PTGS2*^37^ (Fig. 2b-c). Age-downregulated genes were enriched for interferon-alpha response and bile acid metabolism. To distinguish cell-intrinsic aging from compositional change, we repeated the analysis within individual cell types. Most HSPC cell types also showed robust transcriptional aging signatures, arguing against a purely compositional explanation. HSCs alone contained 530 differentially expressed genes, and although overlap in significant genes across cell types was modest, age effects were highly concordant in direction and magnitude, with all HSPC pairwise correlations exceeding 0.5 (Fig. 2d). The strongest age-upregulated and age-downregulated genes within each cell type showed substantial overlap and consistent directional changes (Fig. 2e, Extended Fig. 4a).

The HSPC aging signature was largely sex-invariant, with no genes showing a significant age-by-sex interaction after multiple-testing correction (Extended Fig. 4b). However, this analysis was likely underpowered to detect modest effects, and sex-linked transcripts exhibited some sex-dependent trajectories (Extended Fig. 4c–d). In male donors, Y-linked gene expression declined with age, consistent with a mosaic loss of chromosome Y in aging blood^38^ (Extended Fig. 4e-f).

### Transcriptomic aging follows distinct temporal trajectories

Using a framework to capture non-linear age effects^39^, we identified 3,151 age-associated genes. Clustering their smoothed trajectories revealed four major dynamic modules – two increasing and two decreasing with age – indicating that although HSPC aging appears approximately linear, its underlying molecular programs accelerate at distinct times across adulthood (Fig. 2f). Inflammatory genes showed their steepest increase between ages 35–55, EMT- and cell-cycle-related genes accelerated later around age 50, DNA repair and stress-response homeostasis genes declined earlier around ages 35–45, and metabolic signaling genes declined later around ages 50–65 (Fig. 2g, Extended Fig. 4g-h). This temporal staggering suggests that stress-response decline and inflammatory upregulation emerge earlier in adulthood than cell-cycle, EMT, and metabolic shifts, though longitudinal sampling is needed to rule out cohort effects.

### Single-cell transcriptomic aging clocks capture a robust, intrinsic aging signal

To quantify how strongly age is encoded in HSPC transcriptomic state, we trained aging predictors across our uniformly distributed cohort (Extended Fig. 5). A predictor trained on all HSPCs estimated donor age with a mean absolute error (MAE) of 7.8 years under leave-one-donor-out cross-validation (Fig. 2h). Cell-type-specific predictors showed similar performance overall. Notably, HSC and LMPP predictors had the lowest errors and matched the full-HSPC clock performance despite being trained on fewer cells (Fig. 2i), supporting a cell-intrinsic aging signal across hematopoietic lineages, with limited contribution from changes in cell-type composition. To directly test this, we trained an HSPC compositional clock on 7,768 HSPC-restricted Milo KNN neighborhoods, which showed that, at most, about a third of the aging HSPC signal can be explained by composition changes (R² = 0.26, MAE = 13.8 years; Extended Fig. 5d). Lastly, we applied our HSPC transcriptomic predictor to an external scRNA-seq atlas of 123 donors across adulthood^30^ and observed a strong transfer performance (R² = 0.67, MAE = 8.6 years; Fig. 2j), indicating that this signature is not driven by study-specific batch effects.

### Chromatin accessibility captures a distinct, predictive layer of HSPC aging

To complement the transcriptomic findings and ask whether the same age-associated rewiring was visible at the level of regulatory DNA, we next analyzed the chromatin accessibility modality. Peak-level differential accessibility analysis identified 1,269 peaks that were significantly associated with age in HSPCs (Fig. 3a), including a peak 2.5 kb from *PTGS2* that gained accessibility progressively across the lifespan, mirroring the age-dependent increase in *PTGS2* expression (Fig. 2b; 3b).

**Figure 3:**
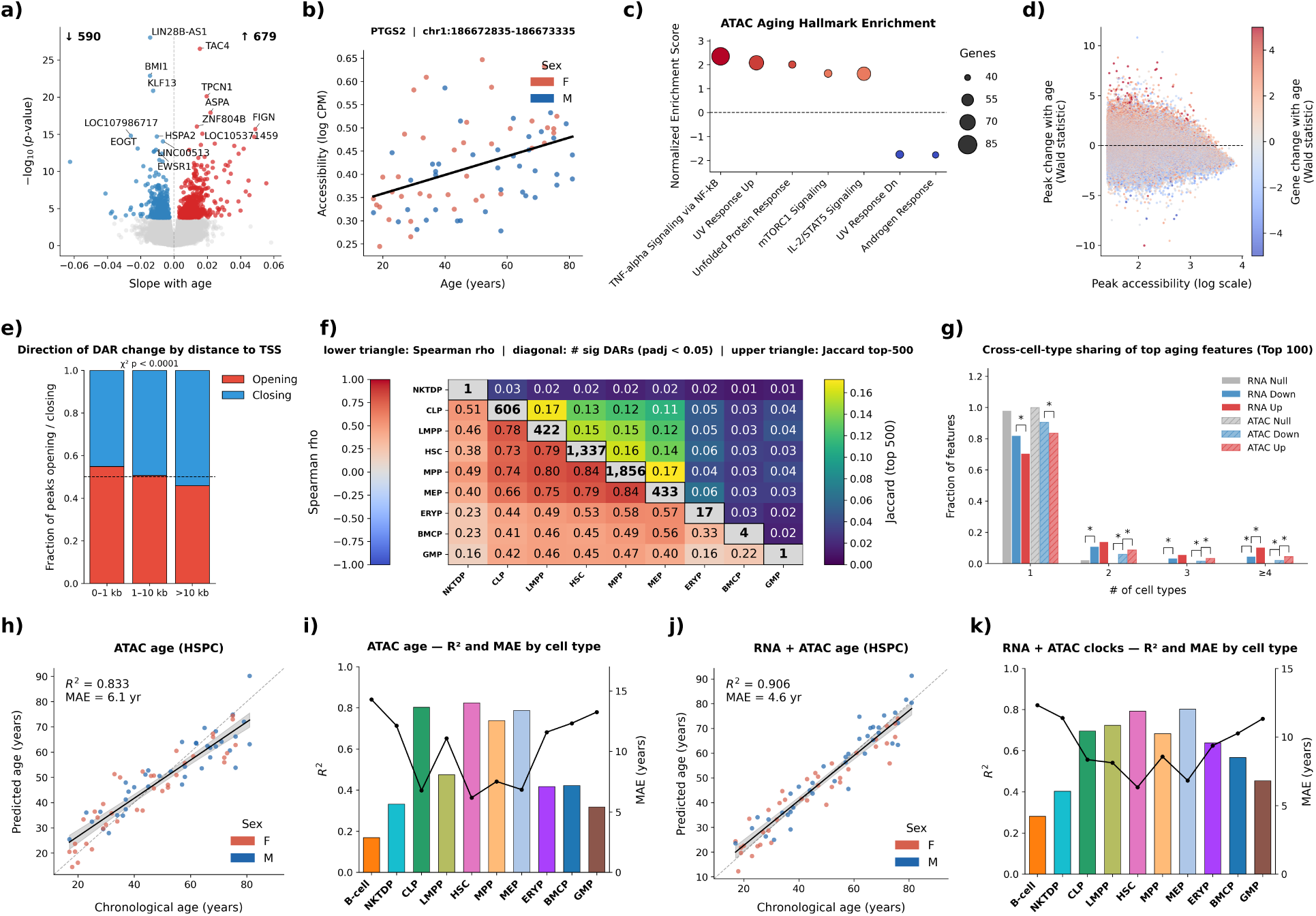
Chromatin accessibility signatures of HSPC aging. **a)** Volcano plot of age-associated chromatin accessibility peaks in HSPCs, annotated by their highest-confidence peak-to-gene linkage. Slope and significance from a pyDESeq2 model with age as a continuous variable and sex as a covariate; 1,269 peaks were differentially accessible with age (FDR < 0.05). **b)** Accessibility of an example age-associated peak linked to PTGS2 (chr1:186672835-186673335) across donors, plotted by donor age and colored by sex. Points show log-CPM-normalized pseudobulk accessibility in HSPCs. **c)** Gene set enrichment analysis of the chromatin accessibility aging signature using MSigDB gene sets, with genes ranked by their strongest age-associated linked peak. Gene sets are ordered by normalized enrichment score; dot size indicates gene set size and color indicates FDR. **d)** Relationship between baseline peak accessibility and age-associated accessibility change. Each point represents a peak, colored by the aging Wald statistic of its linked gene. **e)** The fraction of peaks that increase or decrease with age binned by how far away their assigned gene is. P-value from Chi-squared test. **f)** Similarity of chromatin accessibility aging signatures across HSPC cell types. Diagonal values indicate the number of significant age-associated peaks in each cell type; the lower triangle shows Spearman correlations of peak-level aging statistics across cell types, and the upper triangle shows Jaccard overlap between the top 500 age-associated peaks in each cell type. **g)** Upregulated age-associated features are more broadly shared across cell types than downregulated features. The top 100 aging genes or peaks per cell type are selected and quantified for how often they recur across cell types, plotting the fraction present in ≥ *x* cell types. **h)** Predicted versus chronological age for the HSPC accessibility aging predictor under leave-one-donor-out cross-validation. **i)** ATAC-seq aging predictor performance by cell type. Bar height indicates R^2^, and overlaid lines indicate MAE. **j)** Predicted age compared to chronological age using stacked ATAC+RNA aging predictor. **k)** ATAC+RNA aging predictor performance by cell type.

We linked differential peaks to putative target genes using peak-to-gene correlations, defaulting to nearest-gene assignment when no strongly correlated local gene was detected. The median gene was linked to 10 peaks, although the distribution was highly skewed, with 1.2% of genes linked to more than 100 peaks (Extended Fig. 6a). Genes linked to age-associated peaks were enriched for the same pathways identified in the transcriptomic analysis, including inflammatory and stress-response programs (Fig. 3c), and the direction of accessibility changes matched the direction of expression changes in linked genes. Peaks assigned to age-upregulated genes tended to open, and peaks linked to age-downregulated genes tended to close (Fig. 3d, Extended Fig. 6b). Notably, distal peaks were more likely to close with age than promoter-proximal peaks (Fig. 3e, Extended Fig. 6c–d), consistent with preferential erosion of enhancer-linked regulatory elements.

Accessibility aging signatures were also correlated across HSPC cell types, although more weakly in differentiated populations, consistent with the greater cell type specificity of accessible chromatin landscapes (Fig. 3f). In both RNA and ATAC, age-upregulated features were more broadly shared across cell types than age-downregulated features, suggesting that stress-response programs are broadly redeployed across lineages whereas age-associated declines are more compartment-restricted (Fig. 2e; 3g).

To quantify how strongly age is encoded in the epigenome, we trained accessibility-based aging predictors (Extended Fig. 5). An ATAC-based predictor outperformed the RNA clock, achieving R² = 0.83 and MAE = 6.1 years on HSPC pseudobulk samples, compared with R² = 0.76 and MAE = 7.8 years for RNA, with similarly stronger performance across individual cell types (Fig. 3h–i). Furthermore, a nested ridge stack combining the top-performing RNA and ATAC cell-type-specific predictors achieved superior predictive performance at both the pooled-HSPC level (R² = 0.91, MAE = 4.6 years; Fig. 3j) and within individual cell types (Fig. 3k). These results suggests that the transcriptome and epigenome capture partially complementary dimensions of the aging process, and that paired multiomic profiling provides a higher-resolution representation of cellular aging than either modality alone (Extended Fig. 5).

### Aged HSPCs show expansion of stress and myeloid programs and an erosion of lymphoid and identity programs

The strong, coordinated aging signatures across both modalities motivated us to explicitly model the gene regulatory network (GRN) underlying HSPC aging. Using SCENIC+^40^ to integrate paired RNA and ATAC information and infer TF–gene regulatory links across cell types and donors, we reconstructed a GRN comprising 367 regulons across 146 TFs. The inferred network recapitulated known cell-type-specific TFs and lineage-defining regulators of hematopoiesis (Fig. 4a–b), supporting the validity of the GRN.

**Figure 4:**
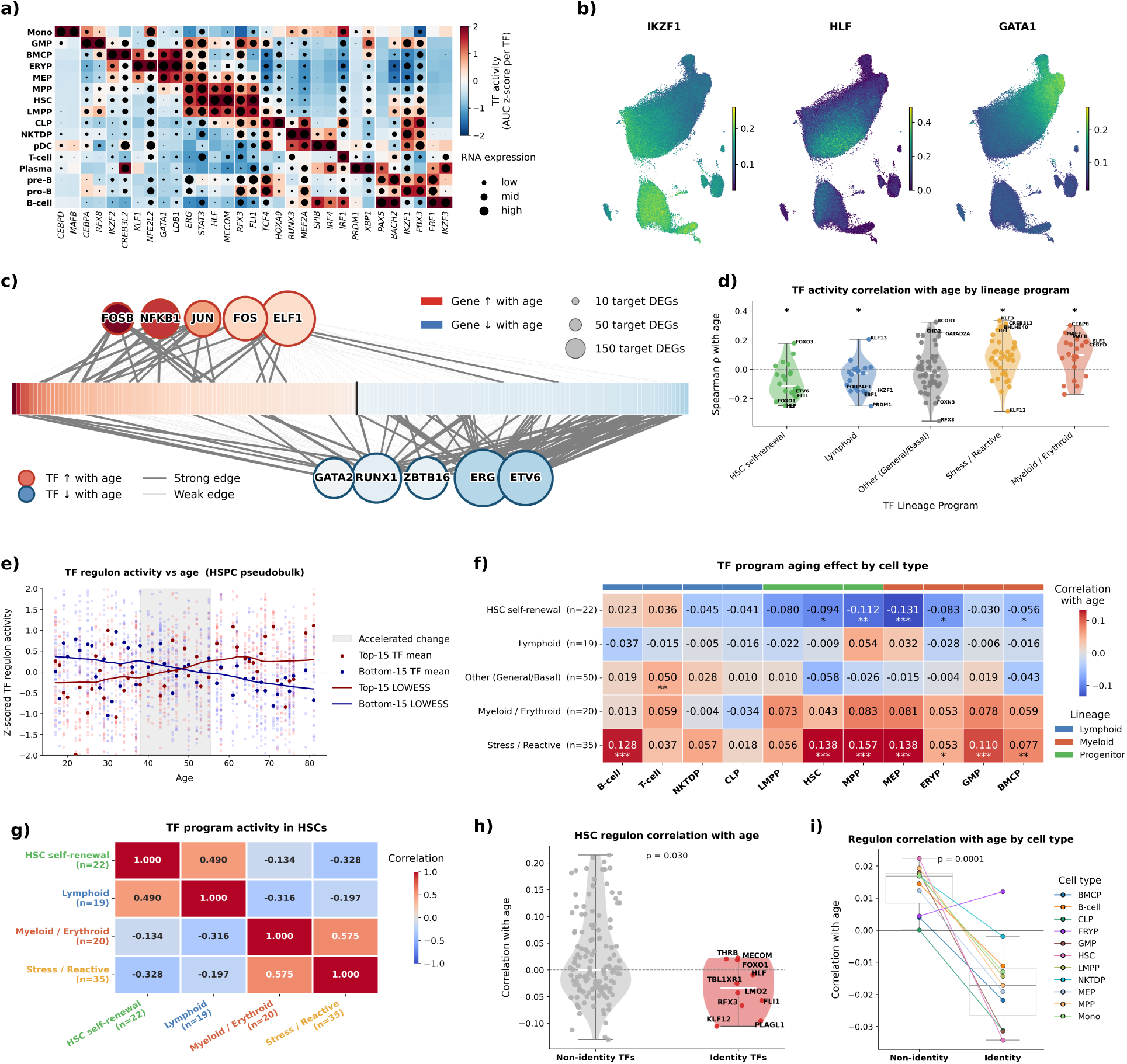
Stress and identity-associated transcription factors underlie age-associated regulatory rewiring in HSPCs. **a)** Cell-type-specific marker regulons from SCENIC+. **b)** Activity of representative master regulators of lymphoid differentiation, HSC self-renewal, and myeloid differentiation overlaid on the ATAC UMAP. **c)** Regulatory network of age-associated differentially expressed genes. The network was restricted to significantly differentially expressed genes represented in the SCENIC+ network and the ten TFs targeting the most differentially expressed genes. The central bar shows target genes ordered and colored by expression change with age. TF nodes are colored by age-associated change in TF activity and size reflects the number of target genes. TF–gene edges with SCENIC+ TF-to-gene importance ≥ 3 are classified as strong edges, with weaker edges shown as faded for clarity. **d)** Age-associated TF activity changes grouped by class: HSC self-renewal, lymphoid, myeloid/erythroid, stress/reactive, or other. Points show per-TF Spearman ρ with age. Asterisks indicate FDR-corrected significance from a one-sample Wilcoxon test of class ρ against zero. **e)** Age-associated TF activity trajectories in HSPCs, showing z-score-normalized activity of the top ten TFs increasing and decreasing with age. **f)** Heatmap of the mean TF activity correlation with age by TF lineage program and cell type. Asterisks indicate FDR-corrected significance from a one-sample Wilcoxon test of class ρ against zero. **g)** Correlation matrix of average TF program activity within HSCs across donors. Per-cell SCENIC+ TF AUC scores were averaged across each donor’s HSCs, z-scored across donors within each TF, and then averaged within each lineage program. Pairwise Pearson correlations between the resulting per-donor lineage scores are shown. **h)** HSC identity-defining TF activity declines with age relative to other TFs in HSCs. HSC identity TFs were defined as the ten TFs whose activity was most specific to HSCs. P-value from Mann–Whitney U test. **i)** Loss of identity-associated TF activity across hematopoietic cell types. Shown is the mean age-associated change in activity of identity TFs in each cell type relative to the average TF activity change in that same cell type. P-value from paired t-test across cell types. *< 0.05, **< 0.01, ***< 0.001, ****< 0.0001.

To identify the regulatory programs most directly associated with aging, we focused on age-associated genes (Fig. 2a) represented in the network and their upstream TFs (Fig. 4c). Genes increasing with age were preferentially targeted by TFs whose activity also increased with age, particularly AP-1 family members and other stress-responsive factors such as NFKB1 and ELF1, whereas genes decreasing with age were predominantly regulated by canonical HSC self-renewal and identity-associated TFs, including GATA2, RUNX1, ERG, and ETV6. Age-increasing TFs also showed stronger and more selective links to age-upregulated targets, whereas age-decreasing TFs were more broadly connected to both up- and downregulated genes (Extended Fig. 6e). This pattern suggests that while stress-associated TFs drive specific age-upregulated programs, the loss of identity-associated TF activity contributes to a broader, passive derepression and destabilization of transcriptional control.

Consistent with this, motif accessibility analyses showed the same overall shift toward stress-responsive regulation and away from core hematopoietic identity programs, with stress-responsive motifs increasing and ETS/GATA motifs decreasing in accessibility with age (Extended Fig. 6f–i). To evaluate these dynamics within a lineage context, we grouped the SCENIC+ age-associated TFs into HSC self-renewal, lymphoid, myeloid/erythroid, stress/reactive, or other classes based on the literature (Supplementary Table 1). These classes showed distinct age-associated behavior: stress/reactive TFs and myeloid/erythroid TFs significantly increased activity with age, while HSC self-renewal TFs and lymphoid TFs declined (Fig. 4d). LOWESS-smoothed trajectories showed that this divergence was greatest between approximately ages 40 and 60, paralleling the midlife acceleration in the transcriptomic trajectory analysis (Fig. 4e).

Notably, these changes were not restricted to the cell types in which the factors normally define identity (Fig. 4f). Within individual cell types, stress and myeloid TF programs increased broadly across both lymphoid and myeloid compartments, whereas HSC self-renewal TFs declined across multiple progenitor states, supporting a cross-lineage, cell-intrinsic regulatory remodeling with age. Across donors, stress and myeloid TF programs co-varied positively within HSCs, whereas HSC self-renewal and lymphoid programs were inversely related to the stress/myeloid axis (Fig. 4g). This suggests that reactive/myeloid expansion and primitive-identity loss are coupled but not perfectly reciprocal features of HSC aging.

### Cell identity transcription factors decline with age

While analyzing TF activity changes by cell type, we observed that canonical cell-identity factors often lost activity with age, even outside the primitive HSC compartment. To formalize this, we defined cell identity TFs as the ten most cell-type-specific TFs in each population and compared their age-associated activity changes with those of all other networked TFs. In HSCs, these identity-defining TFs substantially overlapped the HSC self-renewal class and, as expected, declined with age relative to the rest of the network (Fig. 4h). Strikingly, this pattern extended across nearly the entire HSPC hierarchy: in all cell types, core identity TFs lost activity with age relative to the background TF set, except for ERYP (Fig. 4i). A complementary analysis of motif accessibility showed a similar loss of identity across cell types, consistent with chromatin-level remodeling underlying this regulatory shift (Extended Fig. 6j-k). This pattern is consistent with the age-associated increase in myeloid TF activity, because identity TFs were defined separately within each cell type; thus, even as a shared myeloid program expands across the hierarchy, the TFs most specific to each cell state generally decline with age. This suggests that HSPC aging is accompanied by broad erosion of cell-type-specific regulatory identity across the hematopoietic hierarchy.

### Regulatory entropy increases with age

The broad decline in identity-regulatory programs led us to ask whether aging is accompanied by a more general destabilization of transcriptomic and epigenetic regulation. We use *regulatory entropy* as an umbrella term for loss of organized, reproducible structure in the gene regulatory system, which we quantified here using several orthogonal measurements: cell-to-cell variability, cis-regulatory coherence, chromatin peak sharpness, and donor-specific network architecture (Fig. 5a). Because these measurements capture distinct layers of regulatory organization and have different technical sensitivities, concordant age-associated shifts across them support a common destabilization rather than a single-modality artifact.

**Figure 5:**
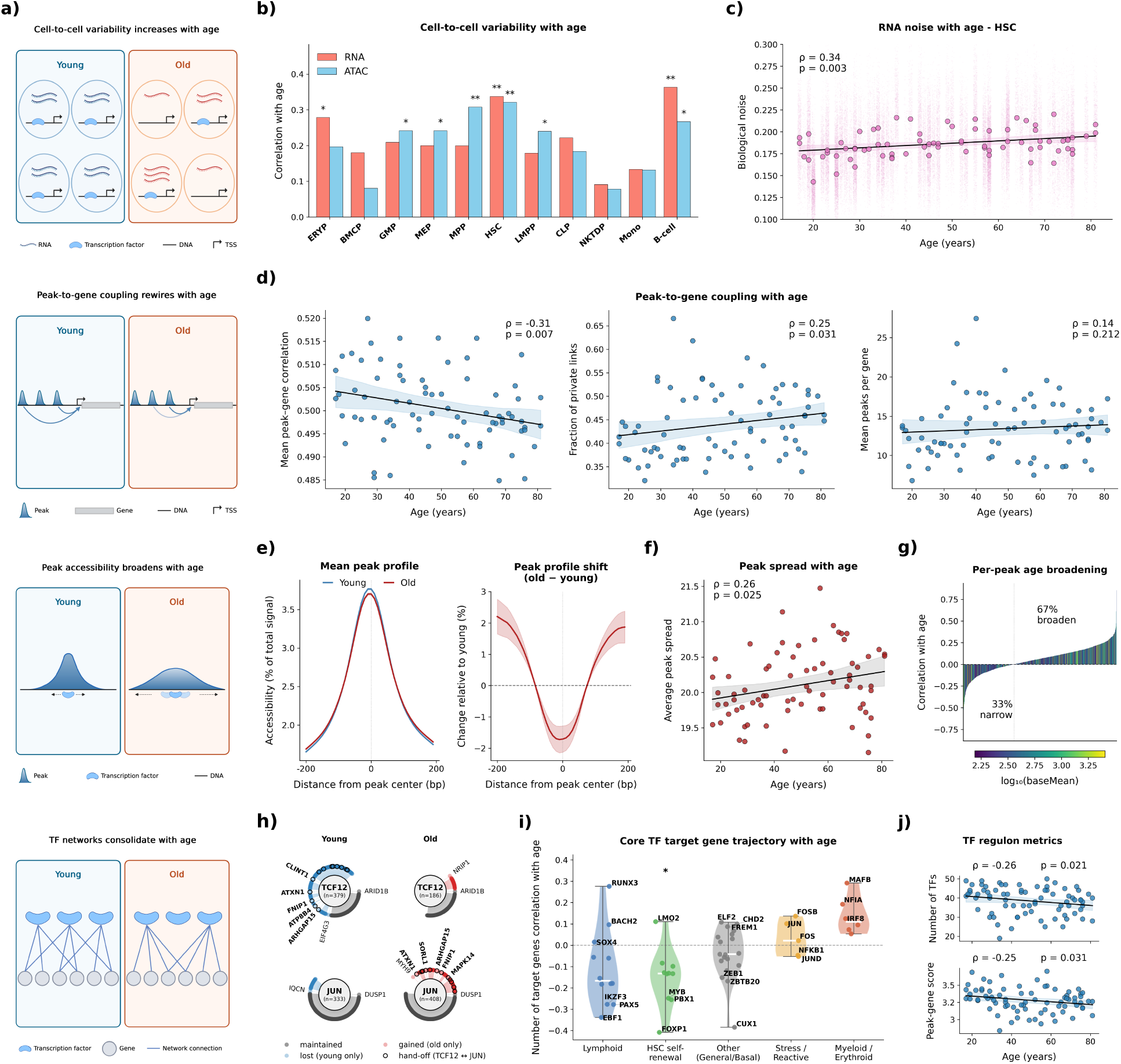
Increasing regulatory instability is associated with age-dependent transcription factor rewiring in HSPCs. **a)** Schematic overview of orthogonal measures of regulatory instability with age. From top to bottom: increase in cell-to-cell variability with age for ATAC and RNA, rewiring and weakening of peak-to-gene coupling, increased peak spread with age, and changes in the TF network. Created using Biorender.com. **b)** Spearman correlation between cell-to-cell variability and age by cell type for both RNA expression and motif accessibility. Asterisks from Spearman p-value. **c)** Scatter plot of HSC biological noise from VarID2 on RNA with age, with large dots representing the mean activity level of all HSC cells by donor, and smaller dots representing individual cells. **d)** Peak-to-gene linkage features across donor age groups. Scatter plots show the mean peak-to-gene correlation, the fraction of rare links (peak-to-gene correlations observed in fewer than 10% of donors), and the total number of peak-to-gene links by donor. **e-g)** Peak-broadening analysis across 5,000 randomly selected peaks with a median log fold-change of zero and an average of at least 150 fragments per donor. **e)** Left, average peak accessibility profiles in young and old donors. Right, difference between old and young profiles. **f)** Scatter plot of average peak spread calculated using FWHM (see Methods) by donor. **g)** Waterfall plot of Spearman correlation between peak spread and age for each peak, colored by that peak’s baseline accessibility. **h)** Representative TFs, JUN and TCF12, that gain and lose target genes with age. Split by young and old, each TF is shown as a center dot connected to its target genes, defined as gene links that appear in at least 50% of samples of that age. Genes and lines are colored by how the TF-gene connection changes with age, with gray representing maintenance, red representing gained edges, and blue edges that were lost. The genes that are lost from TCF12 and gained in JUN with age are outlined. **i)** Correlation between the number of target genes and age, stratified by lineage class. Asterisks indicate FDR-corrected significance from a one-sample Wilcoxon test of the lineage class’s Spearman ρ against zero. **j)** Scatter plots of SCENIC+ network features that change with age. On the top is the number of TF nodes per donor-specific SCENIC+ TF–gene network by age. On the bottom is the average peak-gene importance in a donor network by age. Spearman ρ and p (uncorrected) annotated on each scatter plot panel. *< 0.05, **< 0.01, ***< 0.001, ****< 0.0001.

First, we quantified cell-to-cell variability in RNA expression and found a significant age-associated increase in transcriptomic heterogeneity across HSPC compartments^41^. Extending the same analysis to chromatin, we observed a similar increase in motif-accessibility heterogeneity across cell types, with HSCs showing the clearest and most consistent increase in biological noise across both modalities (Fig. 5b-c).

Second, we examined donor-specific peak-to-gene landscapes as a measure of cis-regulatory coherence. With age, the average strength of peak-to-gene correlations decreased, indicating weaker coupling between accessible regulatory elements and their putative target genes (Fig. 5d). Concurrently, rare peak-to-gene links, defined as correlations observed in fewer than 10% of donors, became more frequent in older donors, even though the total number of links per donor remained stable.

Third, we found evidence of age-associated peak broadening. Across highly covered peaks with near-zero median log-fold change, summit accessibility tended to decrease in older donors while flanking accessibility increased, leading to greater spatial spread of open chromatin (Fig. 5e-f). Although the average effect size was modest, with only a 2% decrease in mean summit accessibility, 67% of peaks showed increased spread with age (Fig. 5g), and peaks with higher baseline accessibility tended to broaden more (Spearman ρ = 0.13, p<0.0001). This broadening is consistent with reduced precision of chromatin accessibility, potentially reflecting less constrained TF binding, nucleosome positioning, or chromatin maintenance with age.

These analyses indicate increased regulatory instability with age, marked by higher transcriptional and chromatin heterogeneity, weaker and less reproducible peak-to-gene coupling, and loss of sharply localized chromatin accessibility.

### Aging rewires and simplifies the regulatory network

To test whether these entropic signatures were reflected in the regulatory network architecture, we inferred SCENIC+ networks separately for each donor. Although donor-specific networks were lower-powered than the combined network, and not all TFs were recovered in every donor, age-associated changes in TF activity tracked changes in the number of target genes more strongly than changes in average TF–target importance, suggesting that aging primarily alters regulon size rather than uniformly strengthening or weakening existing interactions. Consistent with the global network results, stress and myeloid TFs tended to gain targets with age, whereas HSC self-renewal and lymphoid TFs tended to lose them (Fig. 5h–i).

At the network level, the number of TF nodes per donor declined with age, while the total number of genes and edges remained relatively stable (Fig. 5j, Extended Fig. 7c), suggesting that fewer TFs account for a similar amount of inferred transcriptional connectivity in aged donors. Mean region-to-gene importance also declined with age, mirroring the weakening of peak-to-gene coupling described above. These changes indicate a progressive simplification and consolidation of donor-specific regulatory networks with age.

### Baseline regulatory architecture predicts how TFs change with age

We next asked whether intrinsic properties of TF regulons could explain why certain TFs gain activity with age while others collapse. Testing a broad set of regulon-level features revealed several significant predictors of age-associated TF behavior (Fig. 6a; Extended Fig. 7d). TFs that increased with age preferentially regulated GC-rich target promoters and had GC-rich promoters at their own loci. These age-resilient TFs also showed higher basal accessibility and tended to target genes whose associated cis-regulatory regions were more accessible. By contrast, TFs that declined with age were more dependent on distal regulatory elements, reflected by a negative association between TF age slope and the average distance between linked peaks and target-gene transcription start sites. Although these features were strongest at TF target genes, related cis features at the TF locus itself were also modestly predictive (Fig. 6b).

**Figure 6:**
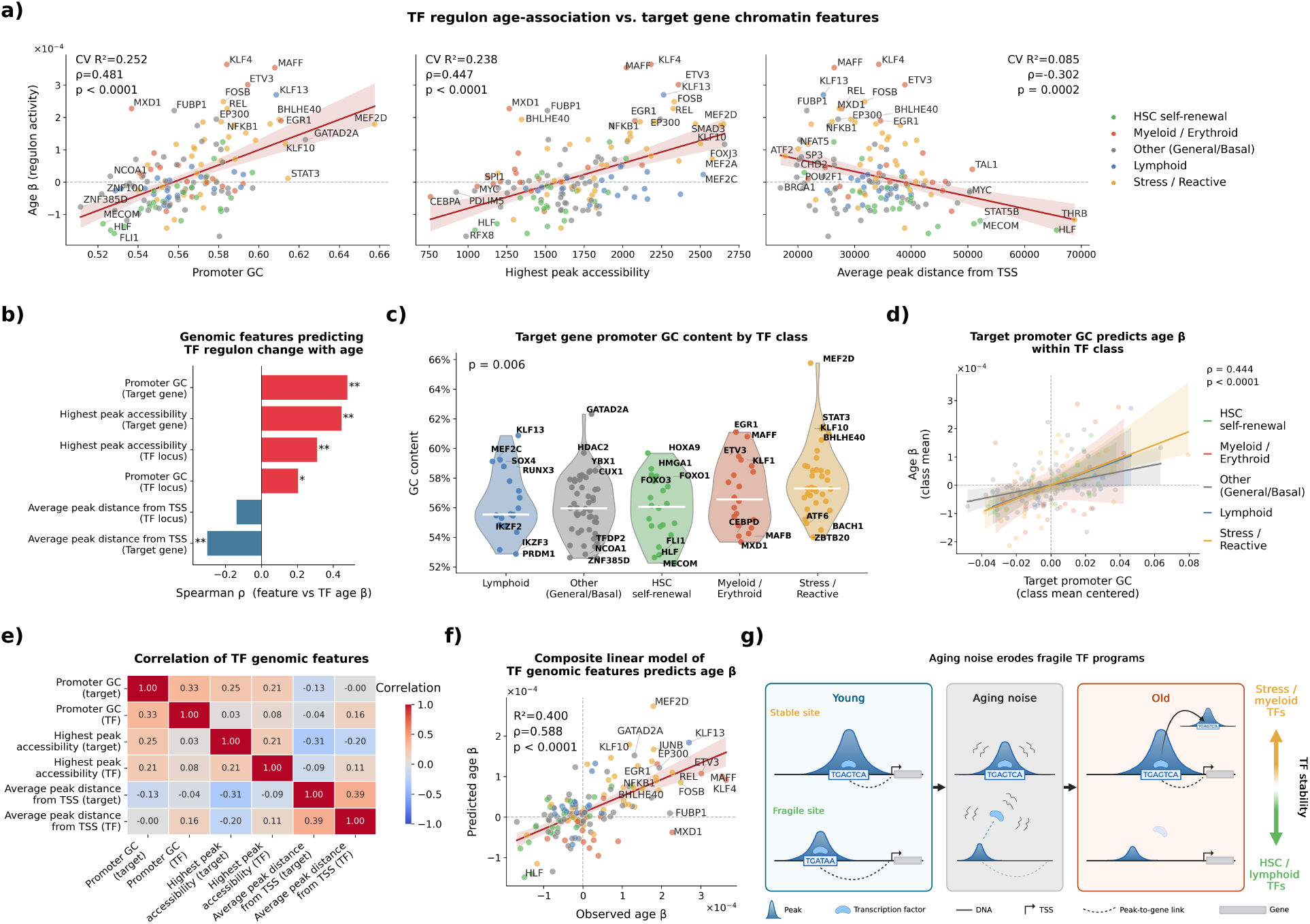
Baseline regulatory architecture predicts TF vulnerability to aging. **a)** Scatter plots of representative regulatory features associated with TF age slope. From left to right: average GC content of TF target gene promoters; average maximal peak accessibility across TF target genes; and average distance of linked peaks from target genes, averaged across each TF’s targets. Points are colored by TF lineage class. Cross-validated R² and Spearman ρ shown on the plot. **b)** Genomic cis (at the TF locus) and trans (TF target genes) features that correlate with a TF’s slope with age. Asterisks indicate FDR-corrected significance from a label-permutation test (1,000 permutations of TF age slopes) on the per-feature Spearman ρ. **c)** Average target-gene promoter GC content per TF, grouped by lineage program. P-value from Kruskal–Wallis test across TF categories. **d)** Spearman correlations between the average target gene promoter GC per TF and that TF’s activity slope with age, split by TF lineage and centered by TF lineage mean change. **e)** Heatmap of Spearman correlations among the six architectural features shown above across TFs. **f)** Elastic Net model trained on the six features above to predict TF age slope under leave-one-out cross-validation. True TF age slopes are shown on the x axis and Elastic Net predictions on the y axis. Cross-validated R² and Spearman ρ shown on the plot. **g)** Schematic model in which increasing regulatory noise preferentially preserves or expands stable TF programs while eroding more fragile identity-associated programs. Created using Biorender.com. * < 0.05, ** < 0.01, *** < 0.001, **** < 0.0001.

Promoter GC content at TF target genes was higher among stress and myeloid/erythroid TFs and lower among HSC self-renewal and lymphoid TFs (Fig. 6c), suggesting that some of the TF-lineage differences might be explained by these baseline TF features. Importantly, the relationship between target-promoter GC content and the TF aging slope persisted within lineage classes (Fig. 6d), indicating that baseline architectural properties, rather than lineage identity alone, shape the magnitude of age-associated TF changes.

Across TFs, these features were only weakly to moderately correlated with one another, suggesting that they capture partially independent properties of regulatory architecture (Fig. 6e; Extended Fig. 7d). Consistent with this, a leave-one-out cross-validated Elastic Net model combining these features predicted age-associated TF activity slope better than any single feature alone (Fig. 6f; Extended Fig. 7e).

To test whether these relationships depended on SCENIC+ inference, we applied an orthogonal TF activity framework based on CollecTRI^42,43^. This independent approach recovered cell-type-specific TF markers and lineage-defining activity patterns (Extended Fig. 8a–b) and again showed increased activity of stress-response and myeloid-associated TFs with age, alongside a relative decline of HSC self-renewal and lymphoid-associated TFs (Extended Fig. 8c–d). It also revealed a significant relationship between basal TF activity and age-associated trajectory, with highly active TFs preferentially preserved or upregulated with age (Extended Fig. 8e–f).

Overall, our findings support a model in which cell-intrinsic HSPC aging can be explained by selective TF vulnerability to aging, where a progressive, entropic destabilization of the gene regulatory architecture preferentially erodes enhancer-driven, identity-associated networks while sparing inherently robust, promoter-proximal stress and myeloid programs (Fig. 6g).

## Discussion

Paired RNA and ATAC profiling across a uniformly distributed adult cohort allowed us to move beyond descriptive signatures of hematopoietic aging and ask how the regulatory architecture is remodeled. This is especially relevant in HSPCs, where myeloid bias, stress activation, and loss of stem-cell function are well established age-related phenotypes whose mechanistic relationship has remained unclear.

Lineage bias in aged hematopoiesis is often interpreted as expansion of myeloid-biased compartments or clones, but our data support a broader model. Although we recover the expected compositional hallmarks of aging, age-associated regulatory signatures persist within individual cell types and across the hematopoietic hierarchy, indicating coordinated cell-intrinsic remodeling. Stress activation, myeloid bias, lymphoid decline, and identity erosion appear to be coupled outputs of a single GRN-level rewiring process rather than independent phenomena.

We propose a framework of architecture-dependent regulatory entropy. Aging increases regulatory noise across the GRN, but each TF program’s vulnerability is shaped by its baseline architecture. Promoter-proximal, GC-rich, highly accessible regulons withstand entropic decay and are preserved or amplified, whereas complex, enhancer-dependent regulons collapse and progressively decline. TF vulnerability thus tracks a regulatory stability axis rather than lineage identity (Fig. 6g).

This model unifies hallmarks of hematopoietic aging that are often treated as independent processes. Stress-responsive and myeloid/erythroid programs, enriched for promoter-proximal architectures, gain activity with age, while lymphoid, HSC self-renewal, and cell-identity programs, enriched for distal-enhancer architectures, decline^44–46^. The age-associated myeloid bias may therefore reflect preferential preservation of myeloid-associated regulatory architecture within individual aging cells, not only expansion of myeloid-biased clones.

The architecture-dependent entropy framework explains the pattern of regulatory change without committing to a specific upstream cause. Promoter-proximal programs may be more resilient because they require less coordinated chromatin regulation, whereas distal enhancer-dependent circuits depend on complex, continuously maintained interactions that are more readily disrupted^47–50^. Multiple processes could elevate regulatory noise, including failure of chromatin-maintenance systems, accumulated DNA damage^51,52^, replication-associated chromatin drift^53,54^, cofactor depletion^55^, and persistent inflammatory signaling. Each would preferentially destabilize the least robust programs, and the model does not require distinguishing among them. Consistent with this, perturbations of chromatin-maintenance machinery in HSPCs produce aging-like phenotypes: EZH2 knockout phenocopies AP-1 overexpression^19^, SWI/SNF loss impairs lineage output^56,57^, H3.3 deletion causes premature exhaustion^58^, and H3K4 methylation depletion causes hematopoietic failure^50^.

This framework also places prior stress-centered models into a broader context. Previous work has proposed that transient stress responses are imperfectly resolved with age, leaving persistent chromatin states that erode cell identity^59–61^. The SIPHON model^19^, for example, proposes that AP-1 overactivation hijacks transcriptional cofactors and closes identity-associated chromatin. Our results extend this view, suggesting that AP-1 activation is one manifestation of broader regulatory instability rather than its primary cause. In an entropic framework, stress gain and identity loss do not need to be causally linked; both can emerge from the same destabilizing process, with stress programs rising because they are architecturally robust and identity programs falling because they are not.

This framework makes several testable predictions. First, regulon architecture should predict aging vulnerability across tissues, not only in hematopoiesis. Cell types whose identity depends on elaborate distal enhancer networks should age faster at the regulatory level than those governed by simpler promoter-proximal circuitry. Consistent with this, disruption of chromatin maintenance in intestinal stem cells selectively erodes distal enhancer programs while preserving promoter-proximal regulation^62^. Second, lineage biases in aging tissues should partly reflect architectural differences among competing fate programs^63^. Third, durable rejuvenation will require suppressing stress TFs alongside restoring identity circuitry, together with the chromatin-maintenance machinery that sustains those programs.

Several limitations should be noted. Our analyses are primarily correlative, and perturbation will be required to establish causal relationships between regulatory entropy, chromatin architecture, and TF aging trajectories. Some conclusions rely on computational GRN inference, which may introduce method-specific biases, though major TF trends are supported by orthogonal approaches. Our cross-sectional cohort also includes limited donor phenotyping beyond age and sex, and unmeasured covariates, such as clonal hematopoiesis status, BMI, and medication use, may contribute to the observed signatures. Finally, while we identify a cell-intrinsic aging signature, we cannot exclude contributions from compositional shifts within annotated cell types, the aging bone marrow niche, or clonal selection, which likely interact with intrinsic regulatory instability.

Our atlas reduces the canonical hallmarks of HSPC aging to a single underlying logic: with age, the regulatory network undergoes a structured, entropic decay. Stress and myeloid programs persist because their promoter-proximal architecture is easy to maintain under regulatory noise, whereas identity and self-renewal programs erode because their distal, enhancer-dependent architecture is difficult to sustain. If this entropic principle generalizes beyond hematopoiesis, it will offer a mechanistic explanation for the age-related fragility of cellular states, stress hyperactivation, and lineage distortion.

## Methods

### Donor recruitment and HSPC sampling

Peripheral blood was obtained from 77 deidentified healthy human donors (39 male, 38 female) through the Rhode Island Blood Center (RIBC) under standard AABB donor eligibility guidelines, which exclude active infection, recent malignancy, and a defined list of medications. No further clinical phenotyping was performed: body mass index, smoking status, comorbidities, medication use, and clonal hematopoiesis status were not assessed. The cohort therefore represents self-identified healthy adults eligible for standard blood donation rather than a deeply phenotyped clinical cohort, and the aging signatures reported here should be interpreted in that context. Samples were obtained under RIBC’s standard donor registration consent framework (RI-FORM-0651), in which donors provide written authorization for research use of donated blood and blood components, and were fully deidentified at source with no donor identifiers transmitted to the investigators. In accordance with Harvard University guidance on human subject research, this work was determined by the investigators not to constitute human subjects research under 45 CFR 46 and did not require IRB review. Donor ages ranged from 17 to 81 years and were distributed approximately uniformly across adulthood, with a target of ten donors per decade and a final per-decade representation of approximately 12 donors per decade except in the youngest (<20 years) and oldest (>80 years) bins. Mononuclear cells were obtained from leukocyte reduction system (LRS) chambers, providing up to ∼1 × 10⁹ starting cells per donor.

Peripheral blood mononuclear cells (PBMCs) were isolated by density gradient centrifugation over Histopaque within 24 hours of collection. Hematopoietic stem and progenitor cells (HSPCs) were then enriched by magnetic-activated cell sorting using the CD34 MicroBead Kit UltraPure, human (130-100-453, Miltenyi Biotec) in MACS buffer, with two sequential LS columns per sample to maximize purity. Enriched HSPCs were counted using trypan blue exclusion on an automated cell counter and cryopreserved in CryoStor® CS10 (07959, Stemcell Technologies) at 200,000–400,000 cells per aliquot. Cells were frozen in CoolCell containers and stored long-term in the vapor phase of liquid nitrogen until use.

### Single-cell multiome library preparation

Cryopreserved CD34⁺ cells were rapidly thawed in a 37 °C water bath for 1–2 min, transferred to a 15 mL conical, and rinsed from the cryovial with 1 mL pre-warmed RPMI + 10% FBS added dropwise. Cells were then sequentially diluted in pre-warmed RPMI + 10% FBS by adding 2, 4, and 8 mL of media at approximately 1 mL per 4 s with 1 min between additions and pelleted at 300 × g for 15 min at room temperature. Pellets were resuspended in residual supernatant, brought up to 10 mL with additional pre-warmed media, and centrifuged again at 300 × g for 15 min. Cells were resuspended in PBS + 0.04% BSA, transferred to RNase-free microcentrifuge tubes with a 0.5 mL rinse of the conical, pelleted at 300 × g for 15 min in a swing-rotor centrifuge, and resuspended in 300 µL PBS + 0.04% BSA. Suspensions were filtered through a pre-wet 40 µm cell strainer and counted on an automated cell counter, then pooled per the multiplexing scheme below into 1.5 mL tubes.

Nuclei were isolated following the 10x Genomics low-input demonstrated protocol (CG000365). Pooled cells were pelleted at 300 × g for 15 min at 4 °C, residual supernatant was removed leaving ∼5 µL, and pellets were lysed in 225 µL chilled lysis buffer (10 mM Tris-HCl pH 7.4, 10 mM NaCl, 3 mM MgCl₂, 1% BSA, 0.1% Tween-20, 0.1% IGEPAL CA-630, 0.01% digitonin, 1 mM DTT, 1 U/µL RNase inhibitor) by gentle pipette mixing. After 3 min on ice, 250 µL chilled wash buffer (10 mM Tris-HCl pH 7.4, 10 mM NaCl, 3 mM MgCl₂, 1% BSA, 0.1% Tween-20, 1 mM DTT, 1 U/µL RNase inhibitor) was added without mixing and tubes were inverted gently. Nuclei were pelleted at 500 × g for 10 min at 4 °C, the supernatant was removed, and the pellet was washed with 400 µL chilled Diluted Nuclei Buffer (1× Nuclei Buffer, 1 mM DTT, 1 U/µL RNase inhibitor) followed by a second centrifugation at 500 × g for 10 min at 4 °C. Nuclei were resuspended in 8 µL chilled Diluted Nuclei Buffer, and 1 µL was used for trypan blue counting on a hemocytometer to assess concentration and integrity prior to transposition.

Transposition, GEM generation, and library construction were performed using the Chromium Next GEM Single Cell Multiome ATAC + Gene Expression Reagent Bundle (1000283, 10x Genomics) following the manufacturer’s protocol (CG000338). Transposition reactions were assembled on ice with 7 µL ATAC Buffer B and 3 µL ATAC Enzyme B per sample, combined with the calculated volume of nuclei suspension and Diluted Nuclei Buffer to a final volume of 15 µL, and incubated at 37 °C for 1 h. Reactions were immediately processed for GEM generation, targeting recovery of 26,000 nuclei per sample.

To increase throughput and reduce batch effects, samples were multiplexed by pooling ∼10,000 cells from one male donor, ∼10,000 cells from one female donor, and ∼5,000 cells from a third donor of either sex per run; donor identity was resolved post-sequencing using genotype-based demultiplexing as described below. Library quality and fragment-size distributions were assessed on an Agilent Bioanalyzer, and libraries were quantified by qPCR prior to sequencing. Libraries were sequenced on a NovaSeq X Plus instrument in 150 bp paired-end configuration, targeting 50,000 read pairs per nucleus for each modality.

### scMultiome processing and preprocessing

Raw sequencing data were obtained from the sequencing provider in BCL format, with one sequencing run per modality. BCL files were converted to FASTQ using bcl2fastq v2.20. FASTQ files from both modalities were then processed with Cell Ranger ARC v2.0.2 to generate paired gene expression and chromatin accessibility count matrices. After generation of raw count matrices and donor demultiplexing with Souporcell v2.0, counts were extracted for each donor and subjected to quality control. ATAC quality control was performed in ArchR v1.0.2 with R v4.4.2, while the remaining downstream analyses were performed in Python v3.13 using Scanpy v1.11 and custom code where necessary.

For RNA, cells were removed if they had fewer than 500 total UMI counts or more than 20,000 total UMI counts, fewer than 500 detected genes or more than 6,000 detected genes, greater than 20% mitochondrial UMIs, or greater than 30% ribosomal UMIs. For ATAC, cells were removed if they had fewer than 1,000 total fragments or more than 75,000 total fragments, fewer than 250 peaks or more than 25,000 peaks, or a transcription start site (TSS) enrichment score below 4 or above 50. TSS enrichment was computed using the ENCODE definition. Cells were additionally filtered based on doublet likelihood. Doublet scores were generated independently using Souporcell for RNA^64^, Scrublet for RNA^65^, and ArchR for ATAC^66^. Cells were retained if they were called singlets by Souporcell and removed if they showed high doublet scores in Scrublet or ArchR based on the empirical score distributions.

### Assigning cells to individuals

Donor assignment was performed using Souporcell v2.0 on the multiplexed data. Cells were clustered according to single nucleotide polymorphism (SNP) profiles, and donor identities were decoded using the expected male:female composition of each pool together with Y chromosome signal and the expected number of cells contributed by each donor. Y chromosome signal was defined for each SNP cluster as a combination of ATAC ChrY peak signal (mean of the per-cell sum of ChrY peak counts) and mean RNA *UTY* expression.

### Dimensionality reduction and clustering

For RNA, 3,000 highly variable genes (HVGs) were identified with scanpy.pp.highly_variable_ genes, after which 9 sex-linked genes were excluded. Raw counts were normalized to 10,000 counts per cell and log1p-transformed. Principal component analysis (PCA) was performed on the HVGs, and the first 50 principal components were retained. A 20-neighbor k-nearest neighbor graph was then computed on these PCs and used for UMAP visualization and Leiden clustering, as implemented in Scanpy. Marker genes for each Leiden cluster were identified in a one-versus-all manner using a two-sided Welch’s t-test implemented in scanpy.tl.rank_genes_groups.

For ATAC, fragments were processed using ArchR’s latent semantic indexing (LSI) workflow. The input consisted of a binary matrix over 500-bp genomic windows spanning the genome, yielding approximately 6 million features. LSI was reduced to 50 dimensions, followed by UMAP visualization and Leiden clustering. To assign biological labels to ATAC clusters, the cells within each ATAC cluster were mapped back to their paired RNA profiles, and marker genes were evaluated using the same iterative annotation strategy as for the RNA clusters. Final cluster annotations were reconciled across modalities as follows: cells with concordant RNA- and ATAC-based labels retained the shared label, and discordant cells defaulted to the ATAC-based label.

Cluster labels were assigned by evaluating the top-ranked differentially expressed genes against curated literature-derived marker gene sets. Marker enrichment was assessed using scanpy.tl.marker_gene_overlap and confirmed via manual gene inspection of canonical markers together with Enrichr-based cell type predictions. Cell type annotations were further refined by reference mapping against the Azimuth bone marrow reference and the Furer et al. circulating HSPC CellTypist model (see “Comparison to external datasets”).

### Comparison to external datasets

We compared our circulating HSPC atlas to three external reference datasets to assess the robustness of the recovered cell states and the relationship between circulating and bone-marrow HSPC compartments.

#### Azimuth bone marrow reference

We mapped our cells onto the Azimuth human bone marrow v2 reference^67^). Azimuth predictions were run using the default reference-query mapping workflow after exporting the scRNA data to Seurat. The per-cell predicted.celltype.l2 label was retained and compared to our annotation as a confusion matrix.

#### Furer et al. circulating HSPC reference

We additionally mapped our cells onto the Furer et al. circulating HSPC atlas using CellTypist with their pre-trained ‘Adult_cHSPCs_Illumina.pkl’ model^30,68^. Query cells were normalized to 10,000 counts per cell and log1p-transformed prior to prediction. Of the model features, 2,245 model features were matched to our gene set. Per-cell labels were assigned by majority voting using the CellTypist default over-clustering resolution of 30 and compared with our annotations as a confusion matrix.

#### Quaranta et al. paired circulating + bone marrow reference

To evaluate how our circulating cells related to bone-marrow-derived HSPCs, we constructed a joint embedding with the Quaranta et al. dataset^27^, which has both circulating and bone-marrow derived HSPCs from the same donors. A 10% random subsample of our cells was concatenated with the Quaranta reference, intersected on shared genes, and jointly processed using 3,000 batch-aware HVGs (batch_key = ‘dataset’), excluding sex-linked genes. The combined matrix was reduced to 50 PCs and batch-corrected with Harmony. In the corrected PC space, each of our cells was assigned a compartment (circulating vs. bone-marrow) by a k = 20-nearest-neighbor vote in the reference, using sklearn.neighbors.KNeighborsClassifier. The fraction of circulating vs. bone-marrow nearest neighbors was compared against the overall composition of the Quaranta reference.

### Cell cycle scoring

Cell-cycle phase was assigned to every cell using scanpy.tl.score_genes_cell_cycle with updated Seurat v3 cell cycle gene lists comprising 43 S-phase genes and 48 G2/M-phase genes^69^. Cells without a confidently elevated S-phase or G2/M score were assigned to G1. The resulting categorical phase labels were used to compute cycling fractions (S or G2/M) by donor and cell type. To compare cell cycle activity across datasets, the same scoring procedure was applied to the Quaranta et al. circulating and bone marrow reference cells. Associations between cycling fraction and age within each cell type were quantified by Spearman rank correlation with Benjamini–Hochberg correction across cell types.

### Lineage regulators of differentiation

To visualize lineage-associated TF dynamics across hematopoiesis (Fig. 1d), HSPC cells were ordered along a simplified lymphoid–HSC–myeloid axis based on smoothed *HLF* expression, similar to prior approaches^30^. For *HLF* and each tested gene, single-cell expression was smoothed over one iteration of the 20-nearest-neighbor RNA graph. Lymphoid-lineage cells were ordered from low to high *HLF* expression, whereas HSC and myeloid-lineage cells were ordered from high to low *HLF* expression, producing a bidirectional axis with primitive HSCs centered between the two branches. Cell type colors matched those used in the UMAPs.

### Cell composition changes with age

To quantify age-associated changes in cell composition, we calculated the relative abundance of each cell type per donor. Associations between cell type proportion and age were assessed using Spearman rank correlation across tested cell types. In Figure 1, proportions were calculated relative to the myeloid HSPC compartment (HSC, MPP, LMPP, MEP, ERYP, GMP, and BMCP), while Extended Fig. 3 includes all cell types. To visualize these shifts non-parametrically on the embedding, we computed UMAP density per donor age group (young ≤35 years, middle 35–60 years, old ≥60 years) using scanpy.tl.embedding_density on the RNA UMAP, after downsampling each age bin to the same number of cells. These same age groups are used throughout the paper when age binning was applied.

### Differential cell-neighborhood abundance

To complement the predefined cell-type-level composition analysis, we performed neighborhood-level differential abundance testing using Milo^34^ from Pertpy v1.0.6, which is robust to the inherent compositionality of proportion data. A joint k = 100 nearest-neighbor graph was computed on the RNA PCs. Cell neighborhoods were sampled at prop = 0.1 of the graph, and per-donor cell counts within each neighborhood were modeled using pyDESeq2 with a continuous age coefficient and sex as a covariate. Neighborhood cell types were annotated by majority-voting, and significance was assessed at a spatial-FDR threshold of 0.01. Per-year log fold-changes were visualized as bee swarm plots and overlaid on the UMAP (Extended Fig. 3d-e).

### Differential gene expression with age

Age-associated differential expression was performed on pseudobulk RNA profiles using pyDESeq2 v0.5.2, with age modeled as a continuous covariate and sex included as a covariate. Analyses were performed both across all HSPCs and within individual cell types by pseudobulking cells by donor. Genes were pre-filtered to those with at least one count in at least 20% of donors. Significant genes were defined at Benjamini–Hochberg FDR < 0.05.

To quantify similarity of age-associated transcriptional signatures across cell types, we computed two complementary metrics for each pair of cell types. First, we calculated the Spearman correlation of Wald statistics (log fold change divided by its standard error) across the union of the top 500 differentially expressed genes ranked by absolute Wald statistic in each cell type. Second, we computed the Jaccard overlap between the corresponding top-500 gene sets. As an additional directional measure of cross-cell-type consistency, we constructed a sign-concordance matrix: for each ordered pair of cell types (A, B), we restricted to genes significant in A (FDR < 0.05 in A) and reported the fraction whose age slope in B had the same sign. To visualize the strongest aging signals per cell type, we took the union of the top three upregulated and top three downregulated genes from each cell type and plotted their Wald statistics across all cell types, marking genes that did not pass pyDESeq2 filtering as missing.

### Age–sex interaction analysis

To test whether age-associated expression changes differ by sex, we refit the pseudobulk differential expression model on pooled HSPCs using the design ∼ Age + Sex + Age:Sex, where Age:Sex represents a continuous-by-binary interaction term (male versus female). We extracted the Wald statistic and FDR-adjusted P value for the interaction coefficient. In parallel, we compared the Wald statistic of each gene between the sex-adjusted model (∼ Age + Sex) and a sex-unadjusted model (∼ Age) to identify genes whose apparent age association was attenuated or amplified by sex composition.

### Sex-linked gene expression and mosaic loss of Y

To assess sex-linked gene expression patterns, the top genes differentially expressed between males and females were z-scored across donors and ordered by donor sex and age. To test for age-associated loss of chromosome Y expression in male donors, we computed Spearman correlations between donor age and z-scored pseudobulk expression of Y-linked genes within males only, with Benjamini–Hochberg correction across the Y-linked gene set. To corroborate this signal at the chromatin level, for each cell we summed ATAC fragments falling within any chrY peak; a cell was called chrY-detectable when this sum exceeded the 99th percentile of female cells (>3 fragments), providing an empirical mapping-noise null that accounts for pseudo-autosomal and repeat-derived leakage. For each male donor we then computed the fraction of cells passing this threshold and tested its association with donor age using Spearman correlation. A coordinated age-dependent decline in Y-linked expression and accessibility was interpreted as consistent with mosaic loss of chromosome Y, as previously reported in aging peripheral blood^70^.

### Gene set enrichment analysis

Pre-ranked gene set enrichment analysis (GSEA) was performed using gseapy.prerank, with genes ranked by the pyDESeq2 Wald statistic for age. Enrichment was evaluated against the MSigDB Hallmark 2020 gene set collection, and FDR-adjusted P values from the permutation test are reported. For chromatin accessibility enrichment, peaks were collapsed to genes by assigning each gene the Wald statistic of its strongest age-associated linked peak (see “Linking peaks to genes”). For trajectory-based analyses, genes in each dCor trajectory cluster were tested for over- representation against the same MSigDB Hallmark 2020 collection using gseapy.enrichr, and the top five terms per cluster were ranked by combined score.

### Aging trajectory analysis

To characterize non-linear age-associated transcriptional trajectories, we used a non-parametric age-association framework based on the distance correlation (dCor) statistic, implemented in the Python dcor v0.6 package^39^. For each gene, dCor values and associated P values were computed from HSPC-level variance-stabilized pseudobulk expression against donor age using dcor.distance_correlation and dcor.independence.distance_correlation_t_test. P values were adjusted across genes using Benjamini–Hochberg correction. Genes were retained if they met either of two criteria: FDR-adjusted dCor P < 0.05 or significance in the DESeq2 age model at FDR < 0.05, yielding 3,151 trajectory-associated genes. Following a previous trajectory inference method^71^, for each retained gene, expression values were ordered by donor age and smoothed across donors using LOWESS (statsmodels.nonparametric.smoothers_lowess.lowess, frac = 0.4, it = 3). Each smoothed trajectory was then z-scored across the age axis. Smoothed, z-scored trajectories were clipped to ±2 and hierarchically clustered using Ward linkage on Euclidean distance into k = 4 trajectory modules. Cluster-level trajectories were defined as the mean smoothed z-score profile across member genes. Functional interpretation of each module was performed by gene ontology enrichment analysis on the corresponding gene sets.

### Trajectory inflection-point detection

To localize age intervals at which cluster-level trajectories accelerated, we computed the numerical derivative of the LOWESS-smoothed cluster-mean profile using numpy.gradient. We then identified contiguous age intervals in which the absolute slope (change in z-score per year) exceeded 0.05 for at least 5 consecutive years. For TF trajectories in Fig. 4g, where effect sizes were smaller, a threshold of 0.01 was used. These intervals were highlighted as shaded regions on the trajectory plots.

### ATAC peak calling

Chromatin accessibility peaks were called with the pycisTopic peak-calling workflow against UCSC hg38 chromosome sizes^40^. ATAC fragments were pseudobulked per donor × cell type, and MACS2 was run on each pseudobulk in paired-end mode (BEDPE input, human effective genome size, shift = 73, ext_size = 146, keep_dup = ‘all’, q < 0.05). The per-pseudobulk narrow peaks were merged into a single 500 bp summit-centered consensus set after excluding regions overlapping the ENCODE hg38 v2 blacklist, yielding 643,355 consensus peaks.

### Linking peaks to genes

Peak-to-gene (P2G) links were inferred using ArchR’s addPeak2GeneLinks workflow based on the correlations between chromatin accessibility and gene expression across KNN-aggregated metacells built on a joint reduced-dimensional embedding. Iterative LSI was run on the peak matrix with two iterations, 25,000 variable features and 30 components (addIterativeLSI), and used as the embedding for both global and donor-specific P2G inference. Default ArchR parameters were used, including maxDist = 250 kb, with no explicit per-peak variability or accessibility pre-filtering. For the global analysis across all donors, we retained links with absolute correlation |r| > 0.2. If no gene within the search window exceeded this threshold, the peak was assigned to the nearest annotated gene by genomic distance using ChIPseeker; this fallback was applied to 83% of peaks. For donor-specific analyses, P2G correlations were computed separately within each donor using the same ArchR pipeline and were retained at two thresholds in parallel: an inclusive cutoff of |r| > 0.2 (for donor-level rare/private-link analyses) and a stricter cutoff of |r| > 0.4 (for donor-level mean-correlation statistics).

### Differential chromatin accessibility with age

Age-associated differential accessibility was performed on donor-level pseudobulk chromatin profiles using pyDESeq2, analogous to the RNA differential expression framework. Peak-count matrices were generated by summing fragment counts across all HSPC cells from the same donor, or from the same donor and cell type for cell type-specific analyses. Peaks were modeled using the design ∼ Age + Sex, with age treated as a continuous covariate. Wald statistics for the age coefficient were used as the per-peak test statistic, and peaks were considered significantly differential at Benjamini–Hochberg FDR < 0.05. Peaks were pre-filtered to those with at least one fragment count in at least 20% of donors, matching the RNA pre-filter. Significant peaks were annotated to genes using the P2G mapping described above. To compare accessibility aging signatures across cell types, we computed the same cross-cell-type metrics used for RNA: Spearman correlation of per-peak Wald statistics across the union of the top 500 age-associated peaks for each pair of cell types, and Jaccard overlap of the corresponding top-500 peak sets. We additionally quantified the fraction of top-100 age-associated peaks and genes per cell type that were shared across increasing numbers of cell types, to assess the relative breadth of age-upregulated versus age-downregulated features.

### Aging Predictors

#### Donor-level profiles and BootstrapCells

We trained independent age-prediction models for each cell type represented by at least 60 donors and at least 20 cells per donor (HSC, MPP, CLP, MEP, LMPP, ERYP, BMCP, GMP, NKTDP and B-cells). For RNA, donor-level pseudobulks were generated by summing transcript counts across cells from the same donor and cell type. Only genes detected in at least 5% of cells within that cell type were retained. BootstrapCells were generated by sampling 100 cells with replacement and summing their counts, with 150 BootstrapCells generated per donor. For ATAC, peak-by-cell fragment counts were binarized, averaged across cells from the same donor to obtain the fraction of cells in which each peak was accessible, and log_2_-transformed.

#### Nested cross-validation and feature selection

All multi-cell-type prediction models were evaluated using leave-one-donor-out cross-validation (LOOCV) across the 77-donor cohort. In each fold, gene or peak filtering, stability selection, and hyperparameter tuning were performed using only the 76 training donors. The held-out donor was not used at any stage of feature selection or model fitting. Within each training fold, stability selection was performed by drawing 100 random 80% subsamples of the training donors and fitting a 5-fold LassoCV model to each subsample. Features selected in at least 50% of subsamples were retained. If fewer than five features met this threshold, the cutoff was relaxed first to 30%, and then, if necessary, to the top 50 features ranked by absolute Spearman correlation with age (Extended Fig. 5a). A final ElasticNet model was then fit on the retained features using 5-fold inner cross-validation at l1_ratio = 0.5 (selected by a preliminary grid search over {0.5, 0.7, 0.9, 0.95, 0.99} on the HSC clock) over the default 100-step λ path. For BootstrapCell clocks, predictions for the held-out donor were averaged across that donor’s BootstrapCells.

#### RNA, ATAC, and multimodal predictors

We trained four classes of per-cell-type predictors within the LOOCV framework described above:

1. **BootstrapCell RNA predictors** used the BootstrapCell representation with scaled log-normalization and dropout filtering (lambda = 75), retaining genes expressed in at least 25% of cells from either young or old donors, and were fit using elastic net with alpha = 0.5.
2. **Stability-selected RNA predictors** replaced the age-stratified dropout filter with an age-unbiased filter, retaining genes with mean CPM > 0.02 and donor-level variance above the cohort median.
3. **ATAC predictors** used donor-level binarized pseudobulks, filtered to peaks with mean accessibility between 2% and 95%, donor-level variance greater than 10^-4^, and capped at the top 15,000 peaks by variance.
4. **Multimodal predictors** concatenated RNA pseudobulk features (log_2_ CPM), ATAC pseudobulk features (log_2_ mean binarized accessibility), and, where available, pathway variability features (see below). Each modality was standardized within the training fold before concatenation, allowing stability selection to operate across modalities. Three multimodal variants were trained per cell type: v5c_RNA+ATAC, v5c_ATAC+PW, and v5c_RNA+ATAC+PW.

Correlations between RNA and ATAC LOOCV residuals within each cell type supported partial complementarity between the two modalities (Extended Fig. 5b).

#### Pathway variability features

For each donor and cell type, we computed the standard deviation of log_2_(CPM + 1) across single cells for each gene, then averaged these per-gene standard deviations across all genes in each gene set to generate one variability score per donor per pathway. Gene sets were drawn from the Hallmark, KEGG, Reactome, and Gene Ontology Biological Process collections in MSigDB, retaining sets with 15–200 annotated genes, of which at least 5 were detected in the data. Missing values were imputed by column-wise medians. This yielded 5,118–5,285 pathway features per cell type (Hallmark, 50; KEGG, 175–182; Reactome, 1,369–1,448; GO:BP, 3,524–3,605), which were used as an additional feature block in multimodal models.

#### Pooled HSPC predictors

As a bulk-like reference that does not split donors by cell type, we constructed three pooled predictors (FullHSPC_RNA, FullHSPC_ATAC, FullHSPC_RNA_ATAC, Extended Fig. 5c) by aggregating all HSPCs into a single donor-level pseudobulk per modality (mean log_2_(CPM + 1) for RNA; log_2_ mean binarised accessibility for ATAC; per-modality-standardized concatenation for the multimodal variant). The same age-unbiased variance filters (top 5,000 genes, top 15,000 peaks) and the same nested LOOCV stability-selection pipeline were applied.

#### Compositional HSPC predictor

To estimate how much of the pooled HSPC signal could be explained by donor-level differences in HSPC composition alone, with no molecular features, we built an aging clock from the per-donor cell distribution across the RNA embedding. We reused the differential-abundance neighborhoods from the Milo analysis (8,305 anchor cells sampled at prop = 0.1, each neighborhood defined as the k = 100 nearest neighbors in the top 50 principal components) and restricted to the 7,768 neighborhoods whose majority cell type was HSPC. For each donor and neighborhood, we recorded the fraction of that donor’s cells captured by the neighborhood, yielding a 77 × 7,768 donor-by-neighborhood frequency matrix, and trained a LOOCV ridge regression (RidgeCV, alpha grid 10⁻⁸ to 10⁵ over 140 log-spaced points, closed-form LOO inner CV per training fold). The compositional clock reached R² = 0.26 and MAE = 13.8 years (Extended Fig. 5c, charcoal bar; Extended Fig. 5d), substantially below the FullHSPC molecular clocks, but well above a constant-mean baseline (MAE = 16.8 years).

#### Cross–cell-type stacking

Although each per-cell-type clock predicted age accurately within its own training cell, generalization to non-HSPC cell types was uneven (Extended Fig. 5e), motivating an ensemble approach. Donor-level predictions from all base clocks were combined into a stacked age estimate via ridge regression, with stacking weights learned by LOOCV (alphas = 50 log-spaced values between 10⁻¹ and 10³; 5-fold inner CV). We report two leakage-free configurations: Global_all, including every base model with a complete prediction vector and no model selection; and Nested_top_k, in which, for each held-out donor, we recomputed each base model’s MAE on the other 76 donors and admitted only models with MAE < k years (k ∈ {7, 8, 10}, Extended Fig. 5g). Because the per-fold MAE filter never uses the held-out donor’s age, the held-out prediction is never used in choosing the models that produce it. The base models contributing most to Nested_top_10, ranked by mean |β| across LOOCV folds, span all three modalities (Extended Fig. 5e).

#### External validation

We assessed generalizability using the independent CD34^+^ HSPC cohort from Furer et al., comprising 123 healthy donors aged 23–91 years and profiled using 10x Chromium with Illumina (90 donors) or Ultima (33 additional donors). Donors profiled on both platforms were assigned to the Illumina dataset. Ensembl gene identifiers were mapped to gene symbols using the dataset’s feature_name annotation, yielding a 20,601-gene overlap with our dataset. External cell types were mapped as follows: HSC_MPP → HSC + MPP, CLP → CLP, MEBEMP-L → MEP, ERYP → ERYP, and BEMP → BMCP (Extended Fig. 2d). For each cell type, we selected concordant genes by computing donor-level Spearman correlations with age independently in each cohort and retaining genes with the same sign and |ρ| > 0.15 in both datasets (636–1,307 genes per cell type). We then generated library-size-matched BootstrapCells, using 189 cells per BootstrapCell in our dataset and 200 in the external dataset, reflecting the observed 1.06-fold median per-cell library-size ratio; 10 BootstrapCells were generated per donor in the external cohort. The combined datasets were jointly log1p (target_sum = 10^4^), zero-variance genes were removed, and batch correction was performed with ComBat using cohort as the batch variable.Internal LOOCV and external transfer MAE were compared by cell type (Extended Fig. 5h). Per-cell-type external predictions were combined using the same LOOCV ridge stacking procedure described above (Extended Fig. 5i). Because both the concordance filter and stacking weights were estimated using external donor ages, this analysis should be interpreted as cohort-adapted transfer rather than fully frozen external validation.

### Motif enrichment and motif accessibility analyses

Motif enrichment in differentially accessible peaks (FDR < 0.05) was first assessed using HOMER v5.1 (findMotifsGenome.pl) with GC-matched non-differentially accessible peaks as background^72^. The top 12 most significant motifs among peaks increasing and decreasing with age are shown in Extended Fig. 6a, colored by FDR and scaled by the fraction of significant peaks containing the motif. To quantify motif accessibility more globally, we ran chromVAR through ArchR with the JASPAR 2020 human vertebrate core motif database (866 motifs)^73^. This produced per-cell motif deviation scores. For age-association analyses at the motif level, we fit ordinary least-squares models of the form deviation ∼ Age + Sex within each HSPC cell type and across pooled HSPCs.

### Reconstructing gene regulatory networks with SCENIC+

Gene regulatory networks were reconstructed using SCENIC+ v1.0, which integrates scRNA-seq and scATAC-seq to infer enhancer-driven regulons (eRegulons) composed of a TF, its target regions, and its target genes^40^. Per-donor count matrices were concatenated, and cells were subsampled to at most 2,000 per donor to limit memory usage and reduce donor-specific cell count imbalance during network inference. Chromatin topics were learned with pycisTopic using Mallet collapsed-Gibbs latent Dirichlet allocation. A 150-topic model was selected from candidate models ranging from 70 to 150 topics based on model likelihood and topic coherence, and the resulting region-topic probabilities were used as cis-regulatory input to SCENIC+. Motif enrichment was performed using the default pycistarget workflow against the v10nr_clust human motif database, and region-to-gene searches were conducted over 1–150 kb around each transcription start site. Final eRegulons were retained at SCENIC+ default thresholds (rho_threshold = 0.05, min_target_genes = 10). For downstream TF-level analyses, per-regulon statistics were aggregated to the TF level by averaging per-cell activity across positive regulons.

### Aging-associated transcription factor activity analyses

TF activity was quantified per cell using SCENIC+ regulon AUC scores. Donor- and cell-type-level pseudobulk TF activity was calculated as the mean per-cell AUC within each donor-by-cell-type group. Age associations were then tested using ordinary least-squares regression of the form TF_activity ∼ Age + Sex, both across pooled HSPCs and within individual cell types, with Benjamini–Hochberg correction across TFs. TFs were manually grouped into five functional categories: HSC self-renewal, lymphoid, myeloid/erythroid, stress/reactive, and other (general or basal), using a curated lookup table (Supplementary Table 1). Assignments were based on established roles in hematopoiesis and stress biology. Category-level summaries report Spearman correlation between pooled-HSPC TF activity and age for each TF, with category-level significance assessed by one-sample Wilcoxon test against zero followed by FDR correction.

To define age-associated regulatory subnetworks, we restricted the inferred SCENIC+ network to significantly differentially expressed HSPC genes (FDR < 0.05) that were also present in the SCENIC+ graph, then selected the ten TFs targeting the largest number of age-associated genes. Edges were classified according to the direction of TF activity change with age and the direction of target-gene expression change and tabulated as a 2 × 2 contingency structure.

### Identity loss with age

To quantify age-associated loss of cell type identity, we first identified identity-defining TFs and motifs for each cell type using a one-versus-rest AUROC specificity score. For each TF or motif, the area under the ROC curve was computed as: AUC = (R₁ − n₁(n₁ + 1) / 2) / (n₁ · n₀), where R₁ is the sum of per-cell TF activity / motif deviation ranks among cells of the target cell type, and n₁, n₀ are the number of cells inside and outside the target cell type, respectively. AUC > 0.5 indicates a feature is more active in the target cell type. Identity TFs and motifs were defined as the top ten features per cell type by AUC, computed across all donors.

Within each cell type, we tested whether identity-defining features declined with age more strongly than the background set of all non-identity TFs or motifs by comparing the distribution of per-feature age slopes between identity and non-identity groups using the Mann–Whitney U test. To summarize across cell types, we computed the mean age slope of identity versus non-identity features within each cell type and compared the two groups using a paired *t*-test across cell types.

### Donor-specific SCENIC+ network analyses

To examine age-associated rewiring of network architecture, we ran SCENIC+ separately for each donor while projecting into the globally learned pycisTopic topic space. For each donor-specific network, we extracted summary statistics from the recovered eRegulons, including the number of distinct TF nodes, mean number of target genes per TF (genes_per_tf), mean number of linked regions per TF (regions_per_tf), mean region-to-gene importance (mean_r2g_importance), and mean eRegulon edge strength (mean_edge_strength). Each statistic was correlated with donor age using Spearman rank correlation, with Benjamini–Hochberg correction across metrics.

For TF-level rewiring analyses, we restricted to core TFs, defined as TFs recovered in donor-specific networks from at least 30 of the 77 donors. This yielded 52 core TFs among 378 TFs recovered across all donor-specific networks. For each core TF, we correlated per-donor regulon size and per-donor TF activity with age using Spearman correlation.

### Noise analyses

#### Cell-to-cell variability through VarID2

To quantify age-associated changes in transcriptomic and epigenetic heterogeneity, we adapted the VarID2 framework (RaceID v2 in R)^41^, which estimates per-cell biological noise by comparing each cell’s profile to a locally constructed pruned nearest-neighbor background. For each cell type represented by more than 2,000 cells, we pruned a 25-nearest-neighbor graph (pruneKnn) using 2,000 variable genes and large = TRUE, recomputed graph-based clustering (graphCluster, p = 0.01, weighted), and estimated per-cell biological noise using compTBNoise (p = 0.01, gamma = 0.5). Neighborhood-level noise was summarized with quantKnn (p = 0.01, minimum neighborhood size = 5), yielding one biological-noise score per cell. Cell-level scores were then averaged within each donor and cell type to generate donor-level mean noise values, which were correlated with donor age using Spearman rank correlation. To extend this framework to chromatin-level heterogeneity, we modified the VarID2 pipeline to work on per-cell ChromVAR motif deviation scores (866 JASPAR motifs × cells) in place of gene expression. All other parameters were identical.

#### Peak broadening

To test whether individual chromatin accessibility peaks became more diffuse with age, we analyzed a panel of high-coverage peaks at donor resolution. Peaks were first filtered to those with pyDESeq2 baseMean ≥ 150 in pooled HSPCs, yielding a high-coverage set suitable for stable per-donor profile estimation. From this set, we drew a balanced random sample of 2,500 peaks with positive age-associated log_2_ fold change and 2,500 peaks with negative age-associated log_2_ fold change, generating a 5,000-peak panel with an approximately zero mean age effect on bulk accessibility by construction. This ensured that age-associated changes in peak shape could be evaluated independently of bulk accessibility shifts. Three donors with fewer than 2,000 recovered HSPCs (956, 1,307, and 1,941 cells) were excluded because their donor-level bigWig profiles were too sparse for stable peak-shape estimation. For each retained donor, we extracted a ±200 bp accessibility profile around each peak center from donor-level bigWig files produced by pycisTopic, using 10-bp bins (40 bins per peak). Each donor-by-peak profile was normalized to sum to one before shape metrics were computed.

Peak spread was quantified for each donor and peak using full width at half maximum (FWHM). Starting from the maximum-accessibility bin, we identified on each side the first bin at or below half the peak maximum and linearly interpolated between adjacent bins to estimate sub-bin half-maximum crossings. FWHM was defined as the distance in bins between the left and right crossings. FWHM could be estimated for 3,094 of the 5,000 panel peaks; the remaining 1,906 peaks were excluded because stable half-maximum crossings could not be identified in most donors. As orthogonal checks, we also computed three additional spread metrics: radial spread, defined as the expected absolute distance from the peak center under the normalized profile; Shannon entropy of the normalized profile; and a flank-to-center ratio, defined as mean accessibility in the outer quartiles of the window divided by mean accessibility in the inner quartiles. Age associations from these alternative metrics were highly correlated with FWHM-based results, so only FWHM is reported for simplicity. For each peak, per-donor spread values were correlated with donor age using Spearman rank correlation to generate per-peak age associations. For donor-level aggregate analyses, the per-donor mean spread across the full panel of peaks was correlated with donor age using Spearman correlation.

#### Peak-gene changes with age

To quantify age-associated changes in cis-regulatory structure, we analyzed peak-to-gene correlations computed separately for each donor (see “Linking peaks to genes”). From these donor-specific peak-to-gene calls, we defined core links as peak-to-gene pairs recovered at the stricter threshold of |r| > 0.4, and general links as peak-to-gene pairs that passed the inclusive threshold of |r| > 0.2 in a given donor. For each donor, we then computed three summary statistics: mean peak-to-gene correlation among core links, fraction of private links (peaks that appear in under 10% of donors at |r| > 0.2) among all links in that donor, and mean number of peaks linked per gene.

### SCENIC+ TF feature analysis

To identify architectural properties of TF regulons associated with age-dependent activity changes, we computed a panel of cis and trans features for each TF from the SCENIC+ network. Feature values were correlated with each TF’s age-associated activity slope using Spearman correlation. A broader feature panel is shown in Extended Fig. 7d; the main text highlights the subset that remained significant after age-permutation testing with Benjamini–Hochberg correction across the full tested feature set. Cis features described properties of the TF locus itself, including promoter GC content, number of peaks linked to the TF promoter, and mean baseline accessibility of those peaks across pooled HSPCs. Trans features described the TF regulon, including mean target-gene promoter GC content, mean baseline accessibility of TF-linked target regions, and mean genomic distance from linked peaks to target-gene transcription start sites.

### CollecTRI-based TF activity inference

As an orthogonal validation of SCENIC+-derived TF activity trends, TF activity was also inferred using decoupler v2.1.1 with the human CollecTRI resource^42,43^. CollecTRI consists of literature-curated TF–target interactions and does not rely on *de novo* network reconstruction or ATAC data from this study. Per-cell TF activity scores were computed using the decoupler.mlm multivariate linear model with default parameters, then pseudobulked by donor and cell type for age-association analyses. Baseline TF activity was estimated from all donors pooled. Age-associated TF change was quantified by donor-level ordinary least-squares regression of the form TF_activity ∼ Age + Sex, and significance was corrected across TFs using Benjamini–Hochberg FDR.

## Data availability

Raw and processed single-cell multiome (paired RNA + ATAC) sequencing data generated in this study will be deposited in the Gene Expression Omnibus (GEO; accession to be assigned) and released upon publication. Processed AnnData objects, peak-by-cell matrices, and per-donor pseudobulk tables used to generate the figures will be made available through the same accession. External datasets analyzed in this study are available under the following accessions: Furer et al. (GSE285943); Quaranta et al. (GSE253485). Custom code for analysis and figure generation is available at <u>github.com/Harlan144/HSPC_scMultiome_Aging_Atlas</u>. The repository is currently private and will be made fully public upon publication.

## Contributions

A.M.P. conceived the project. H.P.S. and A.M.P. designed the study. A.M.P. led donor recruitment, sample procurement, and wet-lab nuclei isolation and library preparation. H.P.S. led the computational analysis, including quality control, cell type annotation, differential expression and accessibility analyses, gene regulatory network inference, regulon feature analysis, and figure generation. A.D.Y. and G.C. designed and trained the transcriptomic, chromatin, and joint RNA + ATAC aging predictors. R.A.G. processed all raw data, led donor demultiplexing, and contributed cluster annotation and quality-control refinement. V.G. provided guidance on aging-biology framing and aging predictor validation. G.M.C., A.M.P., and V.G. supervised the project and acquired funding. H.P.S. and A.M.P. wrote the manuscript with input from all authors. All authors read and approved the final manuscript.

## Acknowledgements

We thank the blood donors who made this study possible, and the Rhode Island Blood Center (RIBC) for facilitating sample collection. We acknowledge BPF Genomics Core Facility at Harvard Medical School for instrument access, Novogene for sequencing support, Juseong Lee for help with library prep, David Chou for advice on HSPC isolation, the Slush laboratory for sharing preliminary data that informed our quality-control approach, and members of the Church and Gladyshev laboratories for helpful discussions and feedback. This work was funded by the Wyss Institute and an NHGRI T32 predoctoral fellowship to H.P.S. (5T32HG002295-23).

---

**Extended Figure 1:**
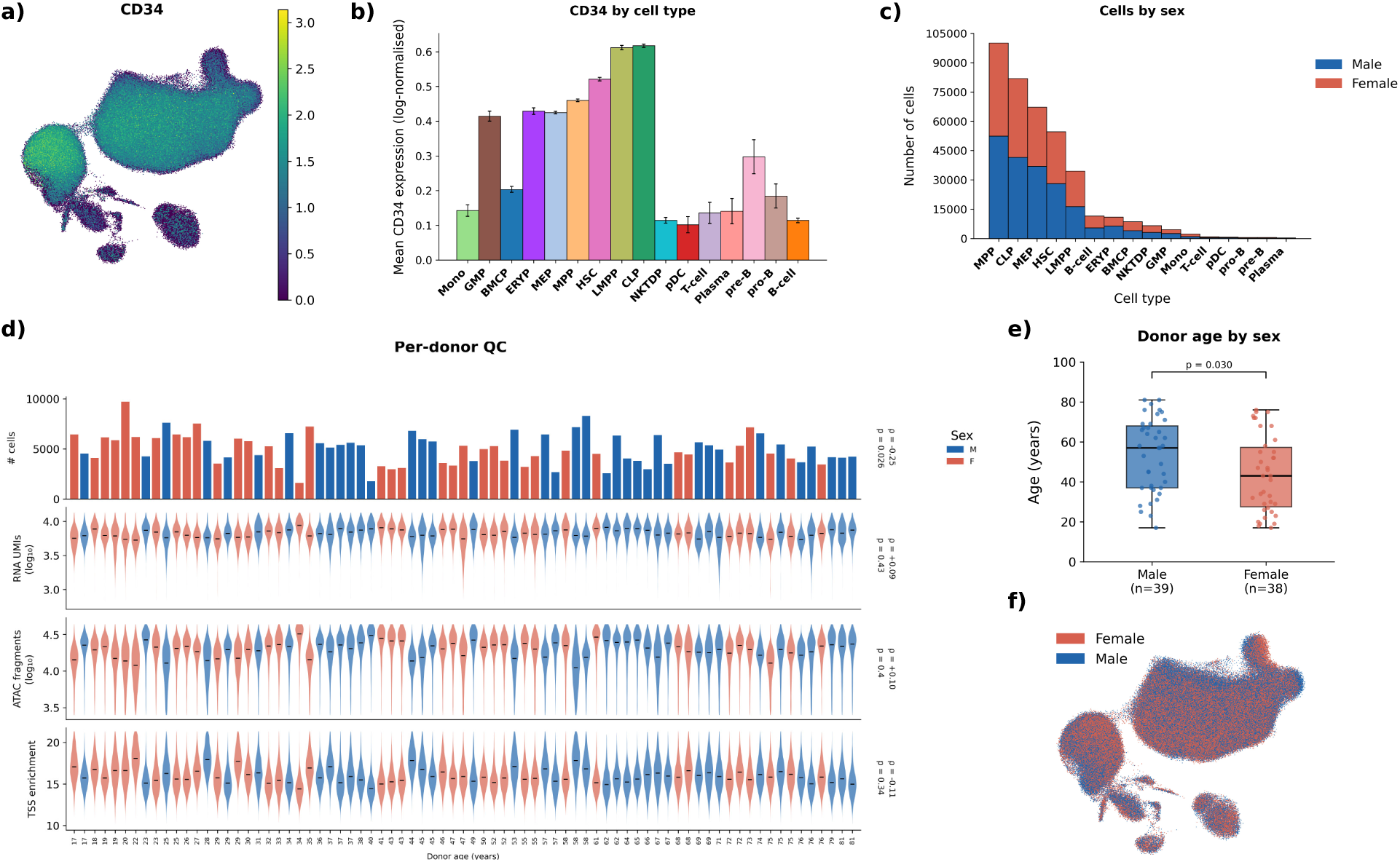
Dataset quality and donor sex-age structure. **a)** CD34 expression across cells projected onto the RNA UMAP. **b)** CD34 expression by cell type. **c)** Number of cells per cell type colored by sex. **d)** Quality control metrics by donors sorted by age and colored by sex. Cell level metrics are shown as violin plots. In order from top to bottom: the number of cells, the log_10_ RNA UMI recovered per cell, log_10_ ATAC fragments recovered per cell, and the TSS enrichment per cell. Spearman ρ and uncorrected p-value annotated per metric. **e)** Boxplot of donor age split by sex. P-value calculated using a two-sided Mann-Whitney U test. **f)** RNA UMAP colored by donor sex.

**Extended Figure 2:**
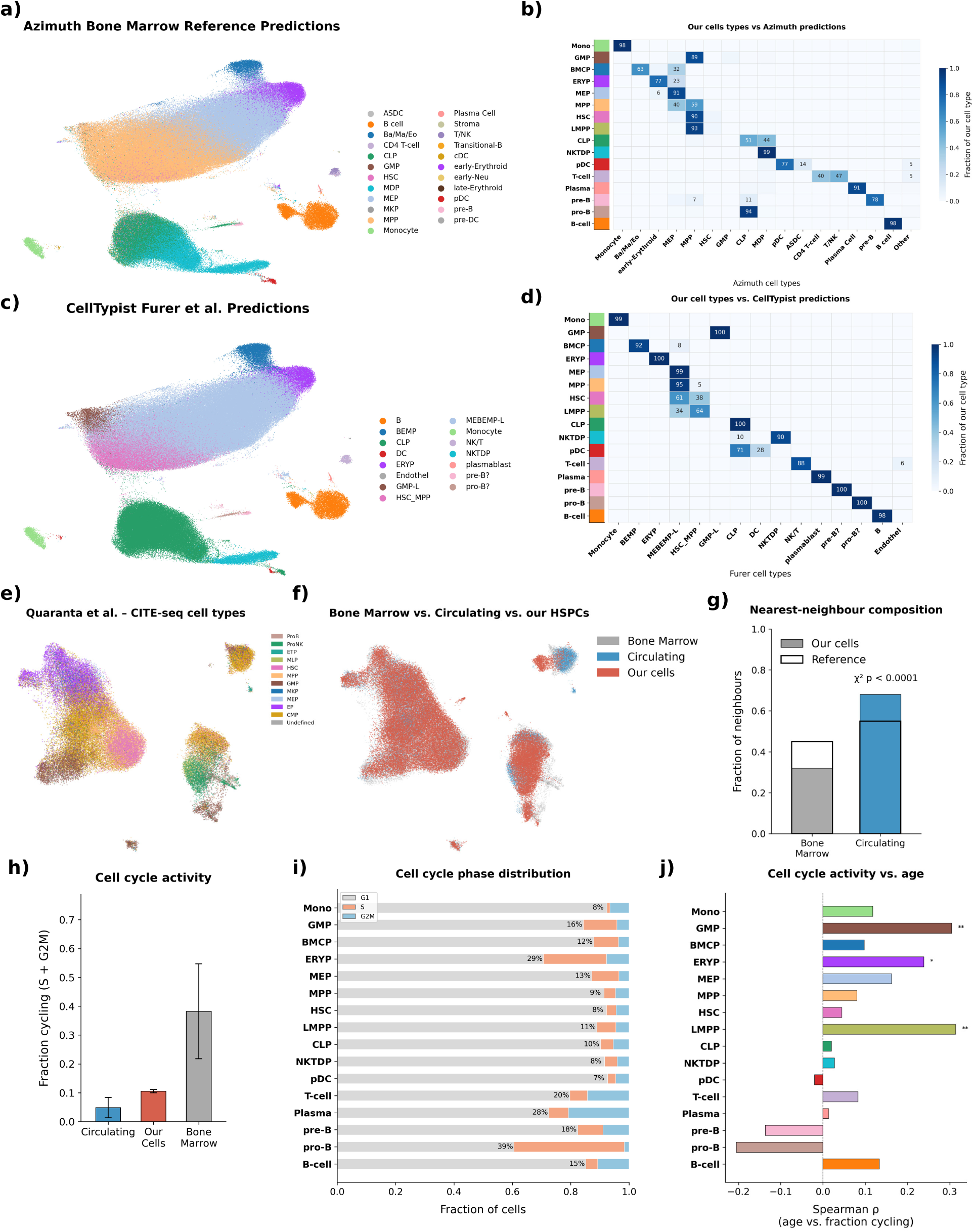
Cross-atlas annotation concordance and cell-cycle profile. **a)** Azimuth Bone Marrow Atlas cell type labels projected on our ATAC UMAP. **b)** Confusion matrix between our annotated cell types and the Azimuth-predicted cell types. **c)** Furer et al. cell type labels from CellTypist projected on our ATAC UMAP. **d)** Confusion matrix between our annotated cell types and the CellTypist-predicted cell types. **e)** Harmonized UMAP of our data (10% random subsample) with Quaranta et al., colored by their cell type annotations. **f)** Harmonized UMAP with colors representing our mapped-on cells, and their cells split by whether they were bone-marrow-derived or circulating. Circulating and Bone-Marrow refer to the two HSPC populations from Quaranta et al., while our cells refer to our HSPC cells. **g)** Barplot showing the fraction of bone-marrow vs. circulating nearest neighbors among the 20-NN of each mapped cell, compared against the overall composition of the Quaranta atlas. P-value from Chi-square goodness-of-fit test. **h)** Barplot of the fraction of HSPC cells that are cycling (predicted S or G2/M phase) by dataset. **i)** Stacked barplots showing the fraction of cells in each cell cycle phase by cell type. **j)** Spearman correlation of the proportion of cycling cells with age per cell type. Asterisks indicate p-value from a per-cell-type Spearman ρ. * < 0.05, ** < 0.01, *** < 0.001, **** < 0.0001.

**Extended Figure 3:**
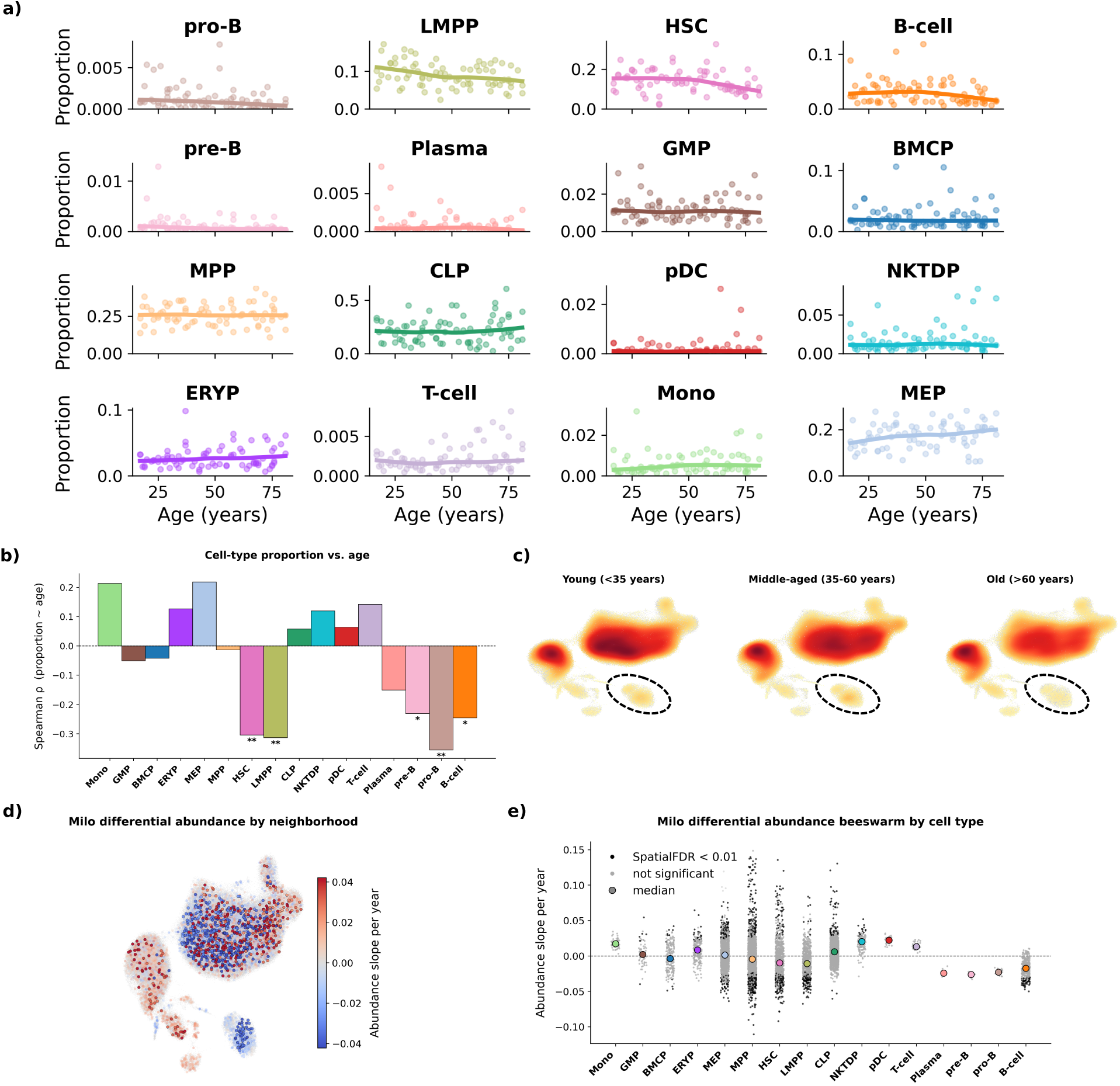
Age-associated compositional shifts across hematopoiesis. **a)** Scatter plot of each cell type proportion change with age, with cell types ordered by their cell-proportion correlation with age. **b)** Barplot of cell-proportion Spearman ρ with age by cell type. Asterisks indicate FDR-corrected significance from per-cell-type Spearman correlations with age. **c)** Density map on the RNA UMAP of the full dataset across donor age groups, showing age-associated shifts in cellular composition, highlighting the decline in B-cells. **d)** Milo-inferred cell-abundance changes with age by cell neighborhood, colored by the slope with age after regressing out sex, with significant (FDR < 0.01) spatial neighborhoods shown with larger dot size. **e)** Bee swarm plot of the slope of each cell neighborhood’s abundance per year, with colored dots representing the median slope per cell type and black dots representing significantly differentially abundant cell neighborhoods with age. * < 0.05, ** < 0.01, *** < 0.001, **** < 0.0001.

**Extended Figure 4:**
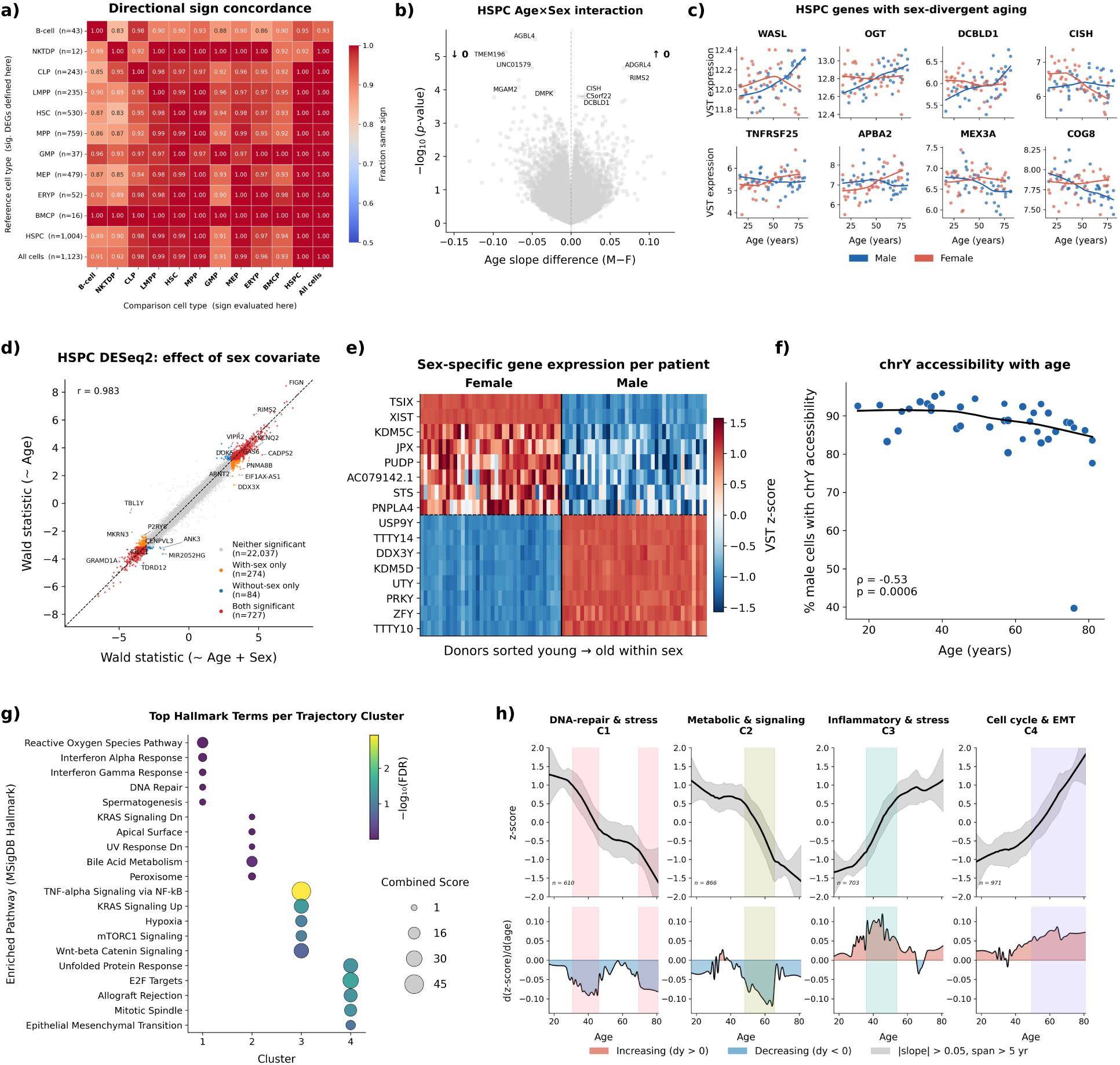
Sex effects and temporal dynamics of the HSPC aging transcriptome. **a)** Directional sign concordance of significant differentially expressed genes between cell types. Rows define the gene set tested; columns show the cell type in which those genes are tested, each cell colored by the fraction of age-associated genes changing in the same direction. **b)** Volcano plot of the Age × Sex interaction term. No gene passes multiple-testing significance (FDR < 0.05). **c)** Expression versus age for the eight age-associated genes showing the largest difference in Spearman correlation with age between sexes. **d)** Scatter plot of age-associated differential expression (Wald statistic per gene) from models fit with and without sex as a covariate. Genes deviating from the diagonal indicate possible sex-biased aging effects. Pearson r annotated on the plot. **e)** Per-donor pseudobulked z-scored expression of sex-linked genes, sorted by assigned sex and age. **f)** Percent of male cells with detectable chrY accessibility versus donor age, consistent with mosaic loss of chromosome Y in older donors. A cell was called chrY-detectable when the sum of ATAC fragments across all chrY peaks exceeded the 99th percentile of female cells (>3 fragments), serving as an empirical mapping-noise null. Spearman ρ and p-value are annotated on the plot. **g)** Top five MSigDB gene ontology terms per trajectory cluster, ranked by combined score and colored by FDR significance. **h)** Smoothed, z-scored trajectory clusters across age (top) and corresponding age-derivative profiles (bottom). Shaded regions indicate intervals of accelerated change, defined as regions where the absolute slope exceeds 0.05.

**Extended Figure 5:**
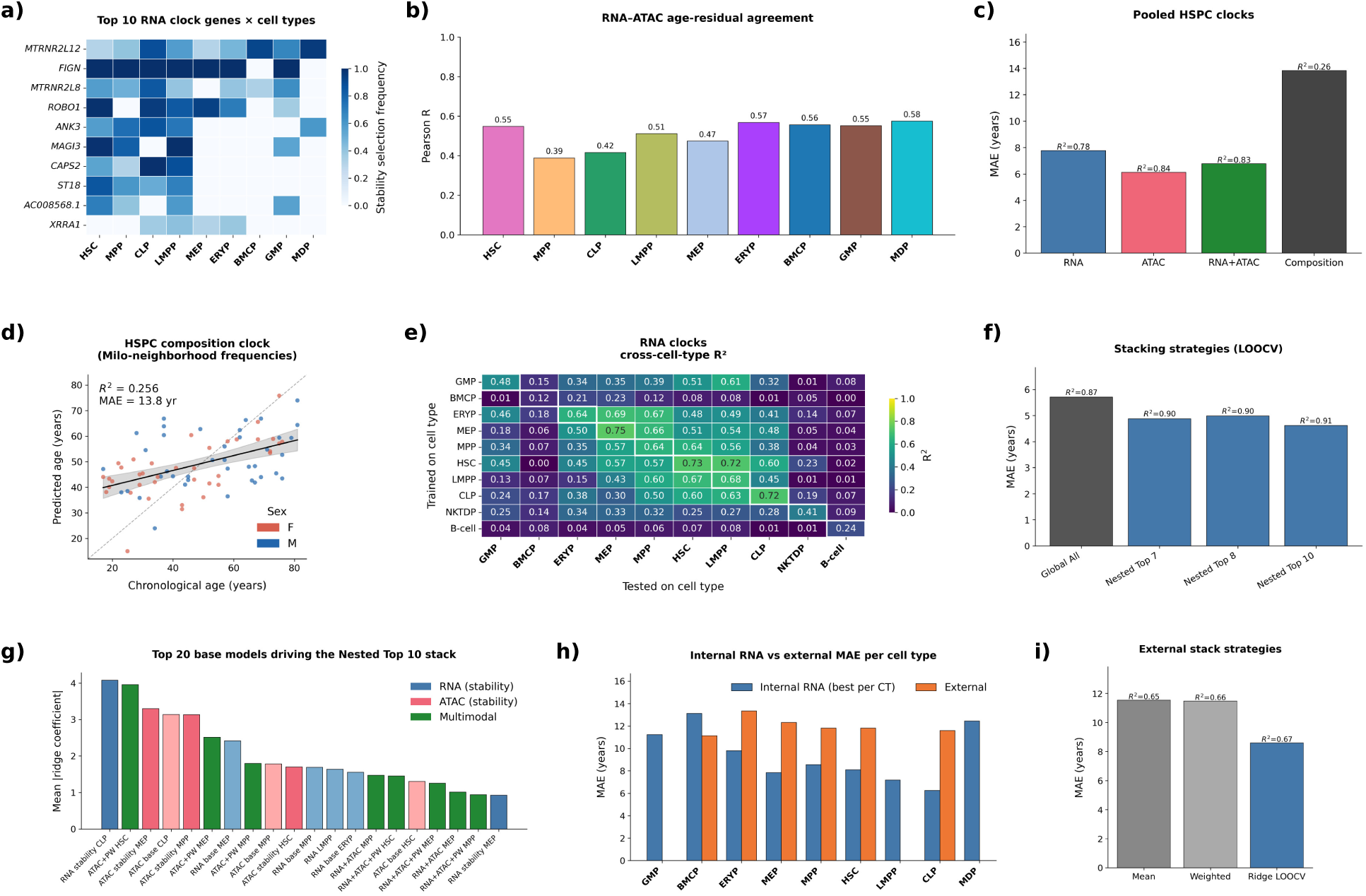
Building transcriptomic and accessibility aging predictors. **a)** Stability heat map of top RNA predictor genes across cell types, showing stability-selection frequency for genes ranked by cross-cell-type coverage and selection frequency. **b)** Per-cell-type Pearson correlation between stability-selected RNA predictor residuals and ATAC predictor residuals in the internal cohort, where residuals are defined as predicted age minus chronological age. This tests whether the two modalities capture shared donor-specific biological-age variation independent of chronological age. **c)** MAE and R² for pooled full-HSPC predictors trained on RNA only, ATAC only, or RNA + ATAC concatenated features, or compositional features without cell type stratification. R² is computed as the squared Pearson correlation between predicted and chronological age. **d)** Predicted vs. chronological age for a composition-only HSPC aging clock trained on per-donor Milo neighborhood frequencies. **e)** Cross-cell-type R² of per-cell-type RNA aging clocks. Each cell shows R² between predicted and chronological age when the clock trained on the row cell type is evaluated on LOOCV held-out donors in the column cell type. **f)** LOOCV MAE by stacking strategy, comparing a flat ridge over all base predictors against nested top-K ridge stacks at K = 7, 8, 10. **g)** Refitted RidgeCV ensemble using the same threshold logic as Nested Top 10 across LOOCV folds, showing the top 20 base clocks ranked by mean absolute standardized ridge coefficient and colored by modality. **h)** Comparison of internal RNA clock MAE and external MAE for each per-cell-type clock. **i)** MAE and R² for the three-cell type stacking strategies on the external cohort: unweighted mean, leave-one-out weighted, and ridge LOOCV.

**Extended Figure 6:**
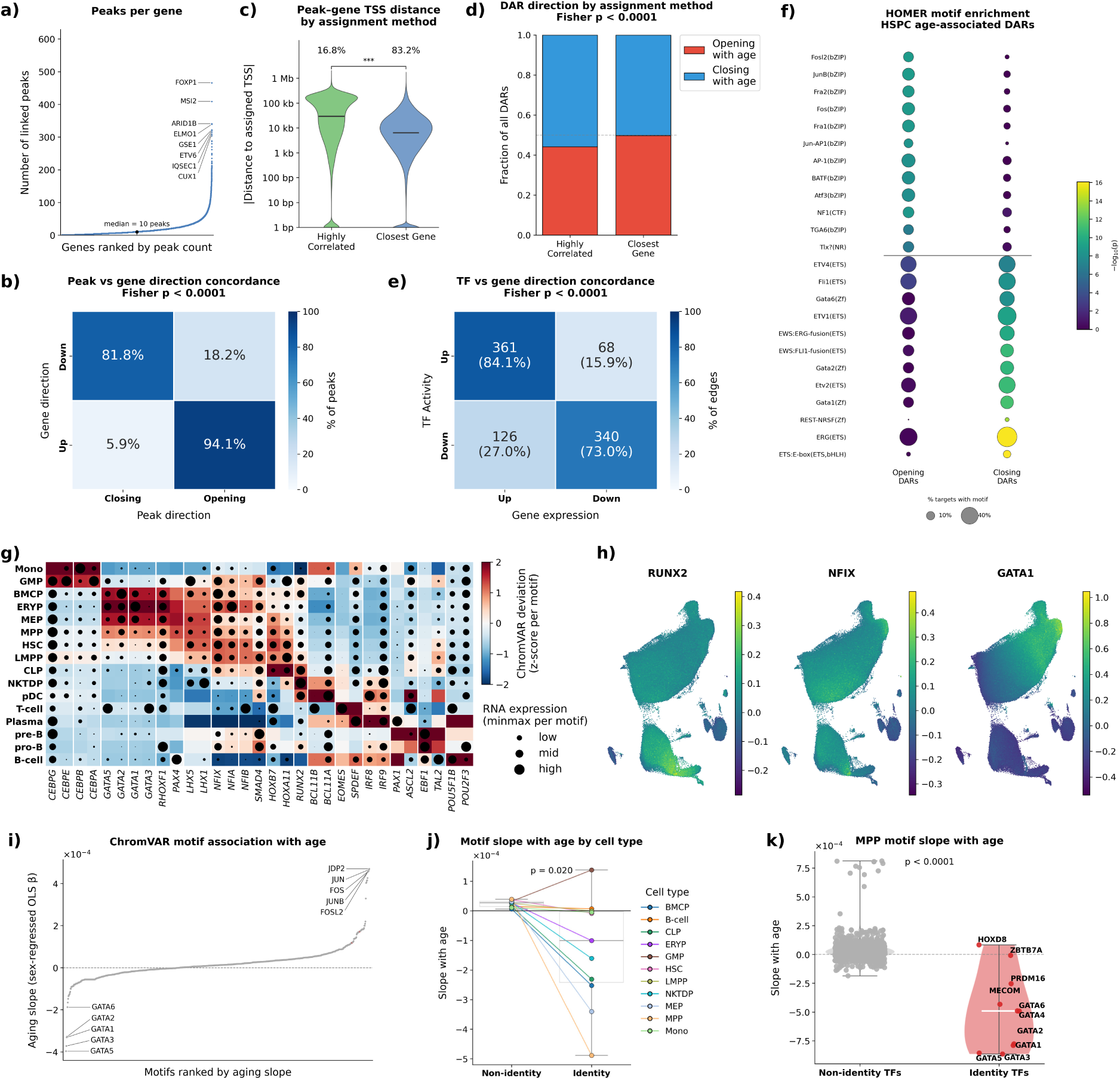
Chromatin and motif-level signatures of HSPC aging. **a)** Genes ranked by the number of assigned peaks. **b)** Contingency plot of significant age-associated peaks grouped by the direction of age change in the linked gene and the peak. Two-sided Fisher exact test. **c)** Distance to transcription start site for peaks assigned by ArchR correlation or nearest-gene proximity. Two-sided Mann–Whitney U test. **d)** Fraction of peaks that open or close with age, stratified by proximity- or correlation-based assignment. Two-sided Fisher exact test. **e)** Contingency plot of TF–target edges grouped by the direction of TF activity and target-gene expression change with age, per Fig. 4c. Two-sided Fisher exact test. **f)** HOMER motif enrichment among significantly up- and downregulated peaks in HSPCs, showing the top 12 motifs per direction. Motifs ordered and colored by FDR significance; point size indicates the fraction of significant peaks containing the motif. **g)** Cell-type-specific motifs from ChromVAR. **h)** Motif deviation scores of representative master regulators of lymphoid differentiation, HSC self-renewal, and myeloid differentiation projected onto the ATAC UMAP. **i)** Waterfall plot of motif accessibility slopes with age. **j)** Loss of identity-associated motifs across hematopoietic cell types, defined as the ten most cell type-specific accessible motifs per cell type. Shown is the mean age-associated accessibility change of identity-defining motifs relative to the average motif change in that cell type. Paired t-test across cell types. **k)** Age-associated accessibility slopes of MPP identity motifs versus all other MPP motifs. Mann–Whitney U test. * < 0.05, ** < 0.01, *** < 0.001, **** < 0.0001.

**Extended Figure 7:**
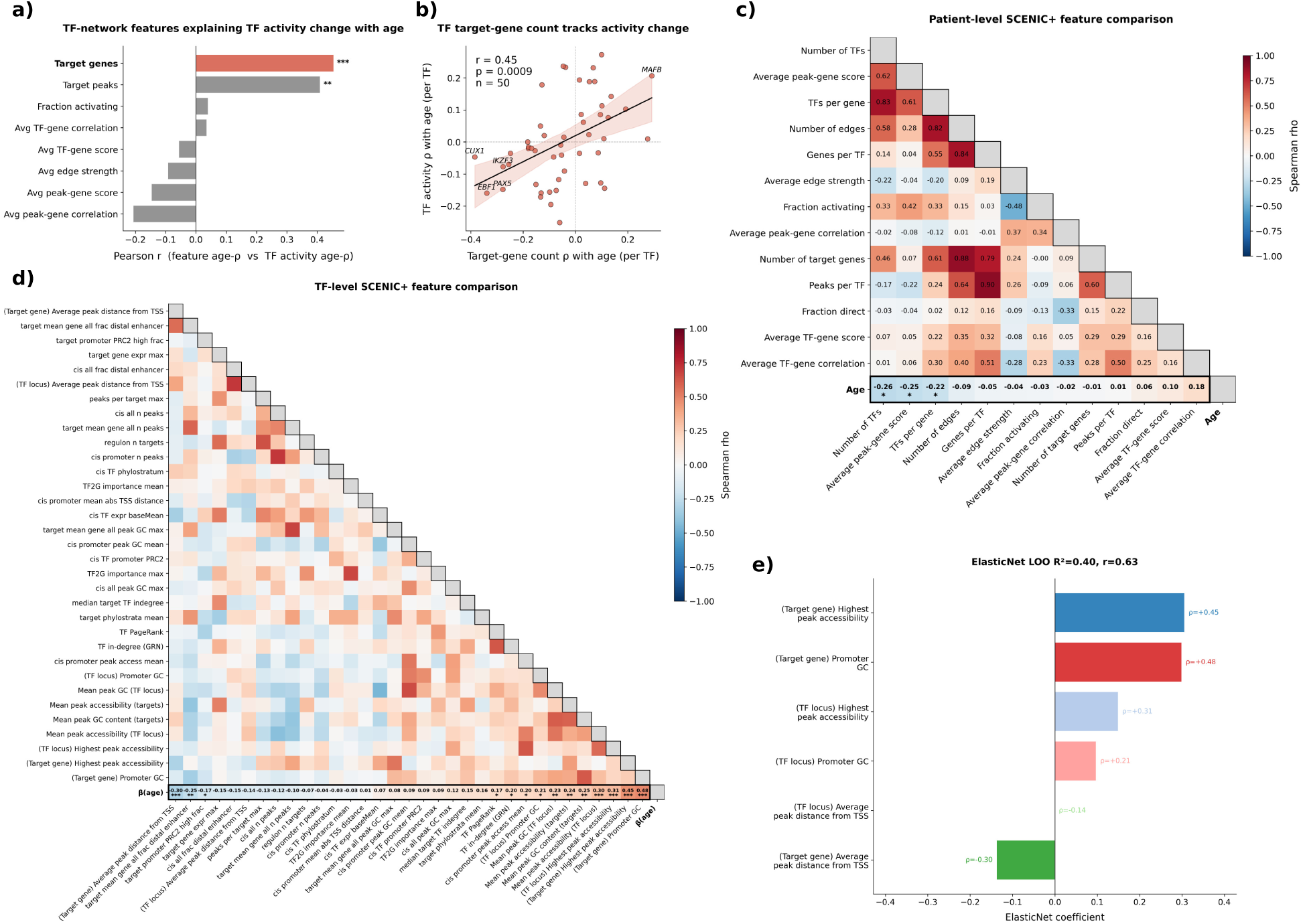
Architectural and network features predicting age-associated TF activity change. **a)** Barplot of TF-level predictors of age-associated TF activity change, comparing TF activity slopes with donor-specific SCENIC+ network features. Only testing TFs recovered in more than 30% of donors. Pearson r between each feature’s per-TF age-ρ and TF activity age-ρ, with asterisks representing FDR-corrected significance. **b)** Scatter plot of the strongest predictive feature: Spearman correlation between donor age and the number of target genes per TF in donor-specific SCENIC+ networks, plotted against that TF’s activity correlation with age in the global network. **c)** Pairwise Spearman correlation heat map of donor-level SCENIC+ network metrics, with correlations against age on the bottom row. Asterisks indicate FDR-corrected significance from Spearman correlation tests. **d)** Correlation structure among TF-level architectural metrics computed from the full cross-donor SCENIC+ network. Correlation between each feature and TF activity slope with age is shown in the bottom row. Asterisks indicate FDR-corrected significance from Spearman correlation tests. **e)** Elastic Net feature weights for the model trained on the six features shown in Fig. 6b to predict TF activity change with age, corresponding to Fig. 6f. * < 0.05, ** < 0.01, *** < 0.001, **** < 0.0001.

**Extended Figure 8:**
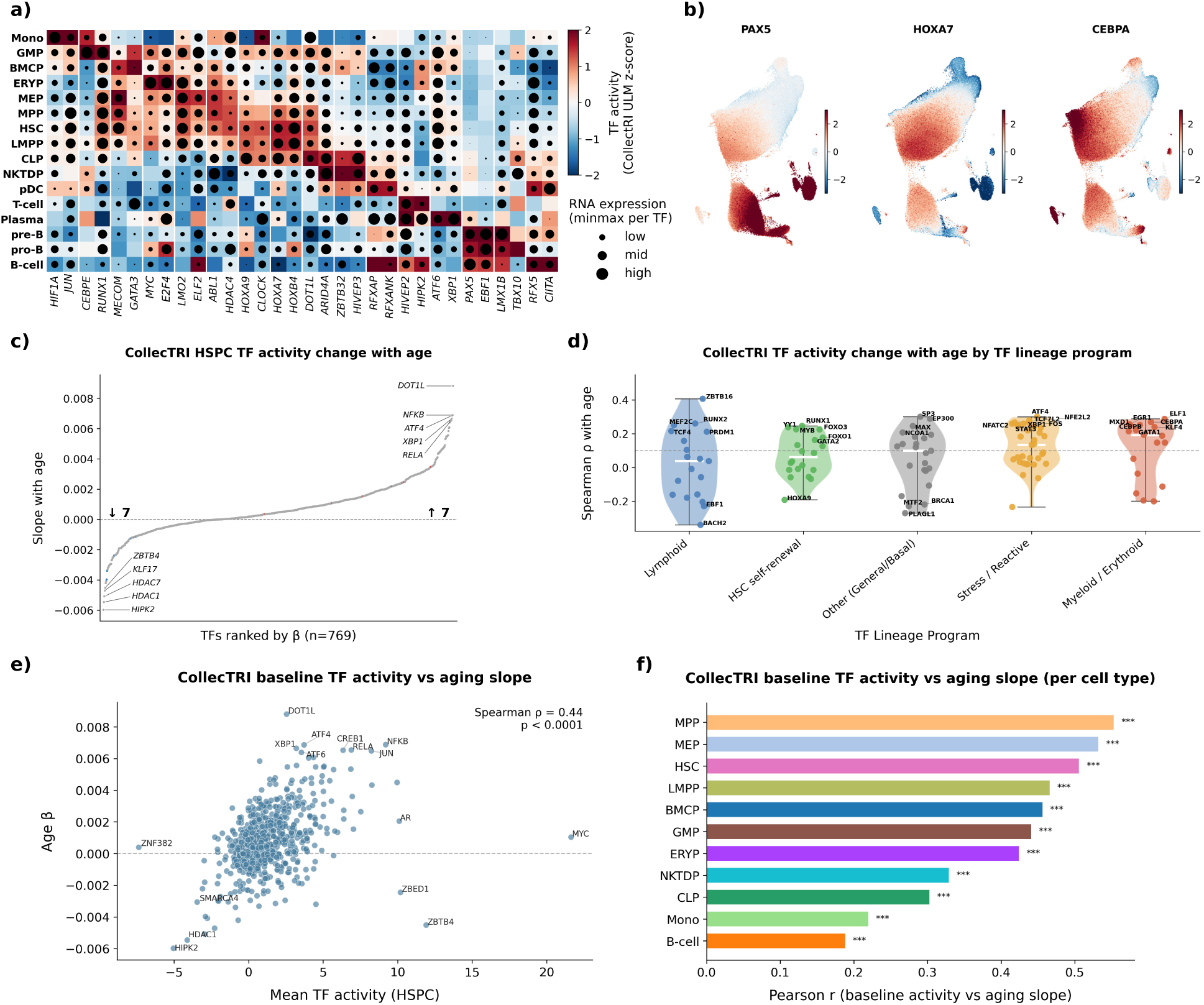
Orthogonal validation of TF aging trajectories with CollecTRI. **a)** Cell type-specific TF activity markers inferred with CollecTRI. **b)** TF activity scores of representative regulators of lymphoid differentiation, HSC self-renewal, and myeloid differentiation projected onto the ATAC UMAP. **c)** Waterfall plot of CollecTRI TF activity slopes with age, after regressing out sex. **d)** CollecTRI TF activity Spearman correlation with age by lineage program. TFs are restricted to those in SCENIC+. The dotted line marks the median Spearman correlation in the Other (general/basal) category. **e)** Scatter plot of mean TF activity across HSPCs versus the TF activity slope with age. **f)** Bar plot of Pearson correlations from the analysis in panel e), stratified by cell type, with CollecTRI activity and age-associated slope calculated separately within each cell type. Asterisks indicate FDR-corrected significance.

